# Patterns of brain-wide associations reflect socioeconomics

**DOI:** 10.64898/2025.12.10.693206

**Authors:** Scott Marek, Meghan Rose Donohue, Nicole Karcher, Caroline Hoyniak, Roselyne J. Chauvin, Ashley C. Meyer, John Miller, Andrew N. Van, Anxu Wang, Noah J. Baden, Vahdeta Suljic, Kristen M. Scheidter, Julia Monk, Forrest I. Whiting, Nadeshka J. Ramirez-Perez, Samuel R. Krimmel, Athanasia Metoki, Sarah E. Paul, Aaron J. Gorelik, Timothy J. Hendrickson, Stephen M. Malone, Rebecca F. Schwarzlose, Carlos Cardenas-Iniguez, Megan Herting, Steven E. Petersen, Joan Luby, Anita C. Randolph, Michael Shanahan, Eric Turkheimer, Benjamin P. Kay, Evan M. Gordon, Timothy O. Laumann, Deanna M. Barch, Damien A. Fair, Brenden Tervo-Clemmens, Nico U.F. Dosenbach

## Abstract

Previous brain-wide association studies (BWAS) cross-sectionally linked a specific behavioral trait, most commonly IQ or psychopathology, to variation in brain function or structure. Here, we expanded the focus of BWAS from effect sizes to interpretability and generalizability by mapping 649 variables to brain function and structure. We compared the resultant BWAS maps to other types of brain data to annotate the BWAS patterns. Socioeconomic status (SES) — not IQ or psychopathology — showed the strongest associations with both resting-state functional connectivity (RSFC) and cortical thickness in the Adolescent Brain Cognitive Development (ABCD) Study. A principal exposome brain pattern, anchored to sensory and motor cortex, captured 34% of the variance across all BWAS maps. This exposome pattern was strongly correlated with the SES and IQ BWAS maps and non-BWAS maps of sleep (EEG), norepinephrine (PET), and stimulants (drug trial), but not cognitive activation maps (task fMRI). Adjusting for SES, reduced brain–IQ associations by 40%. Brain with IQ associations did not generalize, as they could no longer be detected in subsamples drawn from only higher SES backgrounds, while brain with SES associations remained strong in higher-IQ-only subsamples. These findings reveal SES as the principal axis of population-level brain variation, possibly stemming from the sleep deprivation and heightened stress associated with lower SES, since socioeconomics can only indirectly affect the brain.

## Main Text

Brain-wide association studies (BWAS) cross-sectionally link behavioral or environmental variability to measures of brain function or structure (*1–3*) across people. Despite growing sample sizes and increasingly sophisticated analytic approaches (*4–6*), neurophysiological interpretability and generalizability of BWAS associations have remained limited. Recent large population datasets (*7*, *8*) now enable BWAS mapping across the whole phenome and exposome and permit comparison with brain data from other modalities. Identifying patterns that recur across hundreds of variables may reveal shared neurobiological processes, enhancing BWAS interpretability.

Resting-state functional connectivity (RSFC) and cortical thickness are among the most widely used neuroimaging measures of human brain function and structure, respectively. Both are relatively straightforward to acquire, stable within individuals (*9*, *10*) (given sufficient data amount and quality (*11*)) and display measurable inter-individual variability. Spontaneous neural fluctuations (RSFC) are correlated in systematic ways (*12–14*), organizing the brain into approximately a dozen canonical networks based on their functions (*11*, *15*, *16*). Sensory and motor networks show relatively greater day-to-day RSFC variability (*9*, *17*) and sensitivity to stress, arousal (*18*), and drowsiness (*19*). Conversely, the fronto-parietal network (FPN) is more stable day-to-day (*17*, *20*) and supports highest-order abstract cognitive processes (*21*), for example logic and mathematics (*22*, *23*). Effect sizes for cortical thickness are smaller than for RSFC (*1*), but it has been reliably linked with development (*24*), aging (*25*), SES (*26*), and mental health (*26*, *27*).

Prior BWAS typically focused on a single cognitive or clinical variable, most commonly IQ (‘g’; Fig. 1A) or total psychopathology (‘p-factor’; Fig. 1B). However, brain function and structure are known to be affected by extreme environmental exposures, such as childhood abuse (*28–32*) and neglect (*33–35*), institutional care (*36*, *37*), and poverty (*38*), which confer increased risk for psychopathology and cognitive difficulties (*28*, *39*). The extent to which population variability in environmental exposures (e.g., SES), within the typical range, might affect the interpretability and generalizability of common BWAS findings remains unclear. Assessing the relative contribution of brain biology and environmental effects to phenotypic traits is critical for interpreting BWAS.

**Fig. 1.**
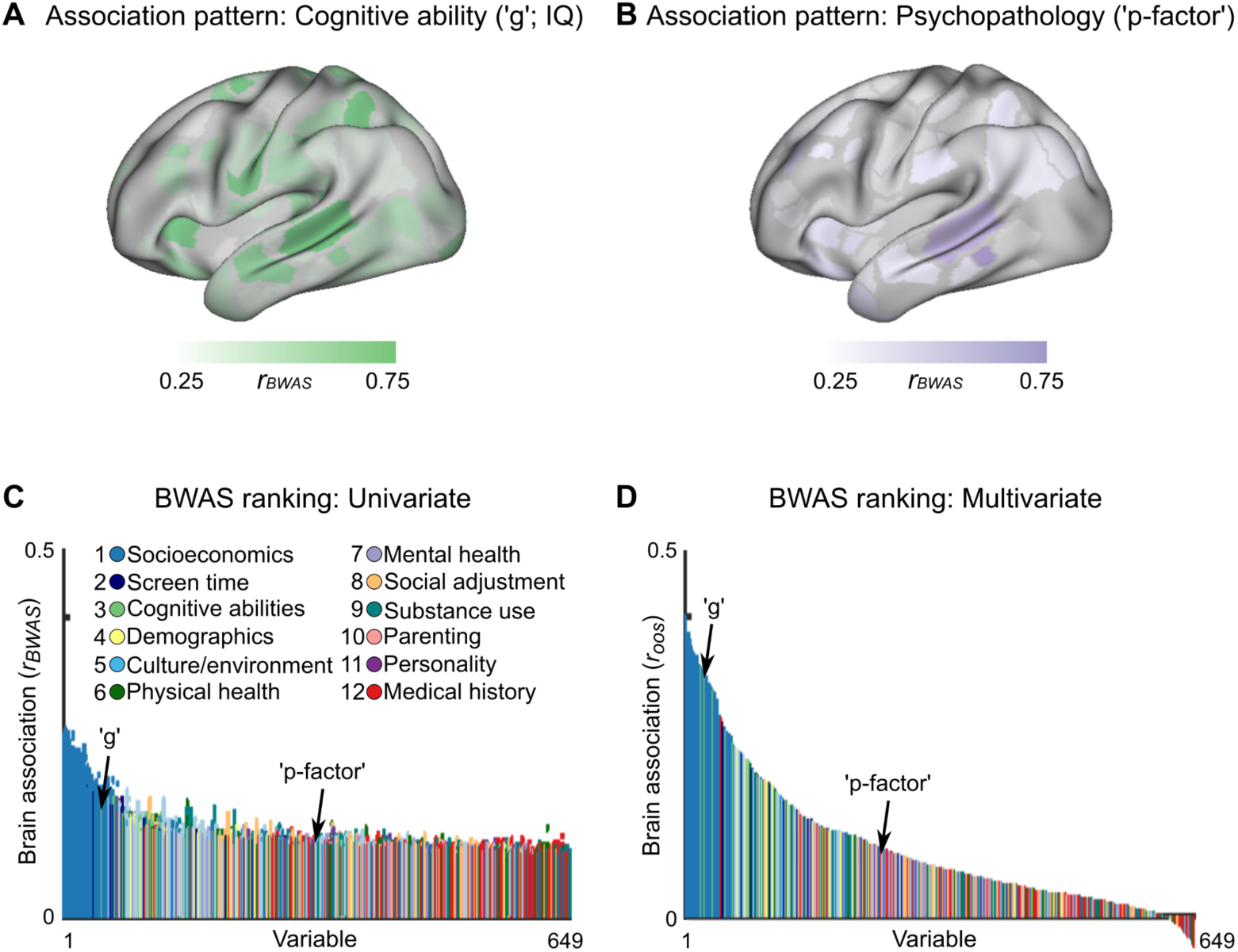
BWAS (brain-wide association study) map strength. Resting state functional connectivity (RSFC) BWAS maps (bivariate correlation, |*r_BWAS_*|; see fig. S1A,B for cortical thickness) for commonly studied exemplar variables: **(A)** cognitive ability (‘g’, IQ scores; NIH Toolbox Cognition Battery, total score) and **(B)** psychopathology (‘p-factor’, Child Behavior Checklist, total score) on a predefined Discovery (*n* = 2,316; training) set of Baseline ABCD data. **(C)** Brain-behavior association strength (y-axis) for 649 non-imaging variables (x-axis). Each colored dot (55,278 dots for each variable) represents a single association between a brain measure (RSFC edge) and a non-imaging variable (see fig. S1C for cortical thickness). Dot color represents the predefined category of the variable. The non-imaging variables (x-axis) are ranked by their 99% confidence interval across all 55,278 associations. All associations were based on at least *n* > 2,000 individuals. **(D)** Multivariate brain-behavior association (bivariate correlation (*r_oos_*) between out-of-sample predicted and observed scores) of brain-based RSFC models for each non-brain variable using ridge regression (see Methods; fig. S1D for cortical thickness). Multivariate brain models were trained using ridge regression on a predefined Discovery (*n* = 2,316; training) set of Baseline ABCD data and subsequently tested on a matched (see Methods) left-out Replication (*n* = 2,263; test) set of Baseline ABCD data. Dot color represents the predefined category of the variable. The non-imaging variables (x-axis) are ranked by their multivariate association strength (*r_oos_*).

Here, we mapped 649 variables (exposome + phenome) to RSFC and cortical thickness in the Adolescent Brain Cognitive Development (ABCD) Study and analyzed the principal dimension of population-level variability. We developed a method for interpreting BWAS patterns, comparative functional pattern analytics, which contrasts BWAS with non-BWAS maps from PET, EEG, drug interventions, and task fMRI. In addition, we tested the generalizability of key brain-wide association patterns in several ways, including through evaluating their relative dependence on subsample characteristics.

## BWAS map strength rankings

The classical phenotypic targets of BWAS, IQ (Fig. 1A; general cognitive ability, ‘g’; fig. S1A for cortical thickness) and total psychopathology (Fig. 1B; ‘p-factor’; fig. S1B for cortical thickness), show distributed brain patterns, with stronger associations for IQ than psychopathology. To cover the breadth of the phenome and exposome, we generated BWAS maps (univariate) for 649 variables in the ABCD Study (Fig. 1C). Ranking all 649 ABCD variables (Supplementary Table 1) by their univariate brain-wide association patterns (*r_BWAS_*) from strongest to weakest revealed socioeconomics, not cognitive abilities or mental health, to be the most strongly associated with brain function and structure (Fig. 1C for RSFC; fig. S1C for cortical thickness; fig. S2 for replication in ABCD Year 2; Supplementary Table 1 for all 649 variables ranked). Of the top 40 variables by association strength with RSFC, 37 were related to a child’s socioeconomic environment, with the remaining three related to sleep and screen time. For cortical thickness, 35 of the top 40 variables were socioeconomic (fig. S1C). Across all 12 predefined categories, socioeconomics demonstrated the strongest brain-wide associations for both RSFC (*F*_(11,648)_ = 51.42, *P* = 6.52 × 10^−81^; one-way ANOVA) and cortical thickness (*F*_(11,648)_ = 20.73, *P* = 3.54 × 10^−36^), followed by screen time and cognition.

The strongest univariate brain-wide association was between the social and economic domain of the Child Opportunity Index (referred to as SES hereafter) and RSFC: *r_BWAS_* = 0.24 (for cortical thickness: *r_BWAS_* = 0.14, 7^th^ strongest). Rankings by effect size (*r_BWAS_*) were highly similar for RSFC and cortical thickness (*r* = 0.79, *P* = 3.01 × 10^−137^) and robust to parameter choices (see Methods). Moreover, rankings replicated in a matched (see Methods), left-out ABCD data set (*40*) (RSFC: *n* = 2,263; *⍴* = 0.83, *P* = 5.22 × 10^−125^; 37/40 top variables were socioeconomic; cortical thickness: *⍴* = 0.64, *P* = 1.47 × 10^−77^; 22/40 top variables were socioeconomic), a larger Baseline ABCD dataset that did not exclude participants for excessive head motion (RSFC: *n* = 7,620; *⍴* = 0.87, *P* = 5.77 × 10^−196^; 37/40 top variables were socioeconomic), and the 2 year follow-up (Year 2) ABCD Study dataset (fig. S1; RSFC: *n* = 2,363, *⍴* = 0.87; 37/40 top variables were socioeconomic). Statistical adjustments for data acquisition parameters (i.e., the use of different MRI scanner types across ABCD sites, inclusion of siblings) did not alter the results (Supplementary Table 1; see Methods).

Multivariate BWAS models yielded stronger associations (*r_oos_*, out-of-sample correlation, between predicted and observed values) for RSFC (Fig. 1D) and cortical thickness (fig. S1D). The maximum multivariate brain association (ridge regression) for SES was *r_oos_* = 0.40 for RSFC (16% of variance explained) and *r_oos_* = 0.36 for cortical thickness. Most of the strongest multivariate associations were with socioeconomic variables (RSFC: 37/40; cortical thickness: 30/40). For RSFC and cortical thickness, the multivariate (*r_oos_*) and univariate (*r_BWAS_*, 99% CI) association strengths were strongly positively correlated (*r* = 0.91 for RSFC and CT, fig. S3), underscoring that their rank order is independent of the analytic approach.

## BWAS map pattern analyses

The strongest brain-wide association map, that of SES with RSFC, was dominated by primary motor and sensory regions (Fig. 2A; see fig. S5A for all brain views), a pattern not observed for cortical thickness (fig. S4A; see fig. S6 for all brain views). The SES motor and sensory BWAS pattern replicated in the adult UK Biobank sample (*n* = 32,572, aged 40-69 years, 95% white British, white Irish, or other White background (*41*); fig. S7), indicating that the socioeconomic brain pattern persists in a homogenous sample with regards to race.

**Fig 2.**
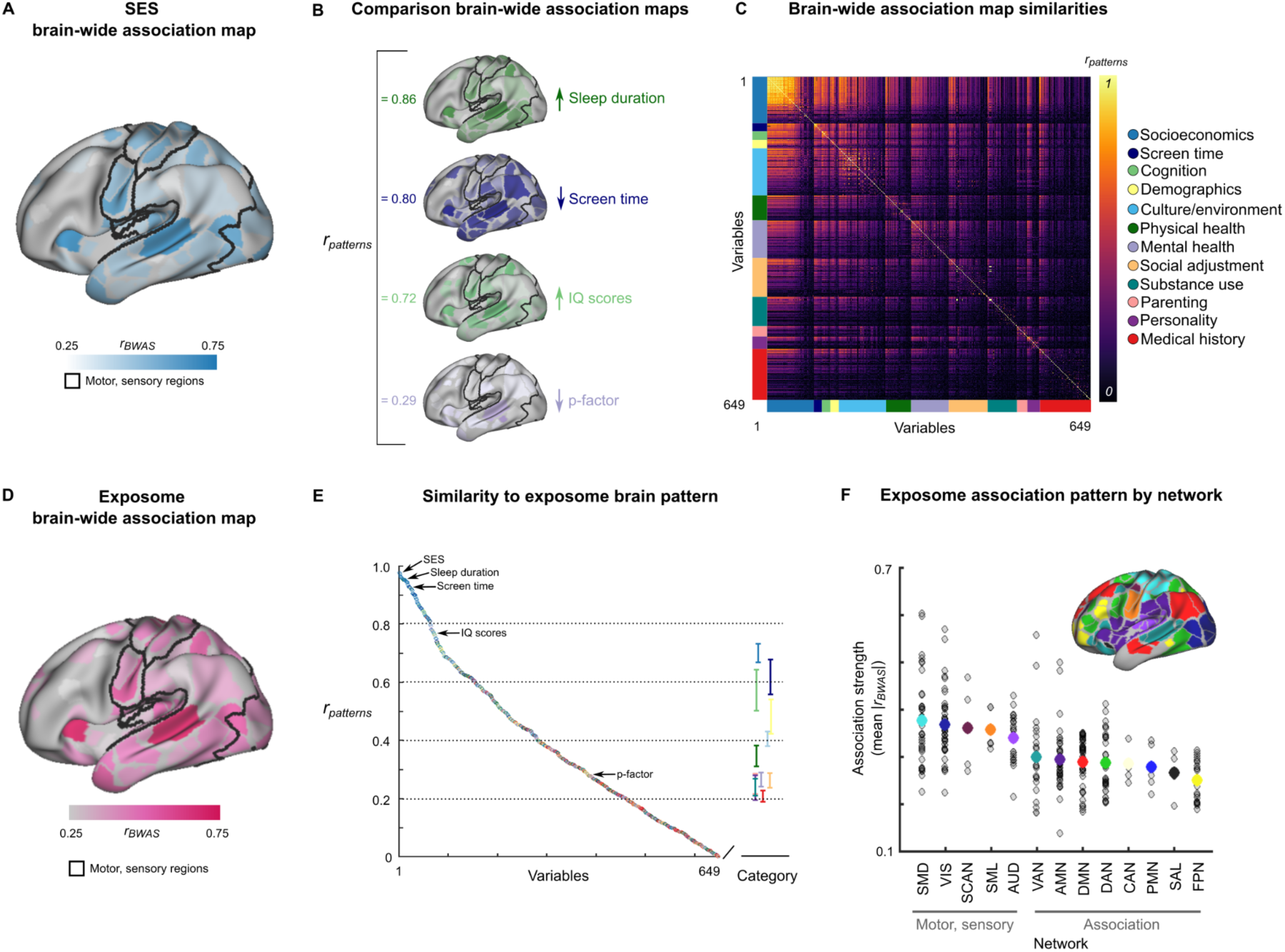
BWAS map pattern analytics. **(A)** Brain-wide association map of resting-state functional connectivity (RSFC) with SES (social and economic domain of the Child Opportunity Index; see fig. S5a for medial wall and right hemisphere). Values are min/max (0-1) normed. **(B)** Exemplar variables for comparison (fig. S5 for additional brain views). The bivariate correlations (*r_patterns_*) between the brain-wide association map for SES (panel **A**) and comparison brain-wide association maps (panel **B**) are listed in the left column of panel **B**. Spin tests revealed significant associations between BWAS maps for SES, sleep duration, screen time, and IQ (NIH Toolbox Cognition Battery, total score, *P* < 0.001, Bonferroni corrected), but not the p-factor. **(C)** Similarity matrix depicting the correlations (*r_patterns_*) for all possible pairs of brain-wide association maps (649 × 649). Variables are organized according to category (right-sided legend) and sorted by association strength within each category. Color tabs on the x- and y-axes reflect the category to which a variable belongs. **(D)** The exposome association map is the first principal component across all 649 brain-wide association maps. See fig. S5 for right hemisphere, medial views. **(E)** Correlations (*r_patterns_*; y-axis) of the brain-wide association maps for each variable (x-axis) with the exposome association map (from panel **E)**. Variables are ranked according to their spatial correlation with the exposome brain-wide association map. The right side of the line break on the x-axis shows the mean +/− the standard error for each category. See fig. S4 for cortical thickness. **(F)** Association strength (mean |*r_BWAS_*|, normed; y-axis) of the exposome map from **D** for each functional brain network (x-axis; see Methods). Each grey circle represents the mean association strength (|*r_BWAS_*|) for a parcel/region. The coloured dot for each network represents the mean association strength across all regions within that network. SMD: sensorimotor dorsal; VIS: visual; SCAN: somato-cognitive action; SML: somatomotor lateral; AUD: auditory; VAN: ventral attention; AMN: action mode; DMN: default mode; DAN: dorsal attention; CAN: context association; PMN: parietal memory; SAL: salience; FPN: frontoparietal.

The BWAS maps for other variables with strong RSFC associations, such as sleep duration (*r_BWAS_* = 0.19; 14^th^), screen time (*r_BWAS_* = 0.17; 32^nd^), and IQ scores (NIH Toolbox Cognition Battery, total score; *r_BWAS_* = 0.16; 59^th^) showed patterns very similar to that of SES (Fig. 2B, correlations to SES BWAS map *r_patterns_*: ↑sleep duration = 0.86, ↓ screen time = 0.80, ↑ IQ scores = 0.72 (all *P* < 0.001, Bonferroni corrected); fig. S4A,B for cortical thickness), while others (e.g., psychopathology) did not (*r_patterns_* = 0.29, *P* > 0.05).

Variables with BWAS patterns similar to SES also had strong direct variable-to-variable correlations with it (fig. S8A; sleep duration *r_vars_* = 0.30, *P* = 1.27 × 10^−50^; screen time *r_vars_* = −0.24, *P* = 2.70 × 10^−31^; IQ scores *r_vars_* = 0.31, P = 6.78 × 10^−53^; see Supplementary Text; fig. S9 for full *r_vars_* correlation matrix). Moreover, BWAS association strengths (fig. S8B; y-axis: *r_BWAS_* 99% CI for each variable) and correlations with SES (fig. S8B; x-axis: *r_vars_*) were strongly related (RSFC: *r* = 0.87, *P* = 6.72 × 10^−198^ cortical thickness *r* = 0.70, *P* = 2.89 × 10^−56^; see fig. S10 for cortical thickness). This relationship suggests that some brain-behavior associations (for example, IQ) are stronger than others (for example, psychopathology) because of their stronger variable-to-variable correlation with SES.That is, IQ has a stronger association with the brain than psychopathology because SES is more strongly associated with IQ than psychopathology (fig. S8A; IQ *r_vars_* = 0.31; psychopathology *r_vars_* = −0.09).

To systematically compare brain patterns across all variables, we correlated the 649 BWAS maps with each other (Fig. 2C; |*r_patterns_*| for RSFC; see fig. S4C for cortical thickness), revealing strong spatial correlations across many socioeconomic variables (bright colors in top left of Fig. 2C for RSFC; fig. S4C for cortical thickness). For RSFC, almost all screen time and cognition BWAS maps were strongly correlated with the SES pattern (|*r_patterns_*| > 0.50; Fig. 2C). Across all non-socioeconomic RSFC BWAS maps, 22% were correlated at |*r_patterns_*| > 0.50 with the pattern for SES (Fig. 2C). While 3% of all non-socioeconomic cortical thickness BWAS maps were correlated at |*r_patterns_*| > 0.50 with the SES pattern (fig. S4C).

The strong neuroanatomical similarity between many BWAS maps suggested a common association pattern may exist across many variables. Thus, we extracted the first principal component (PC) of all BWAS maps (see Methods). The resulting RSFC exposome association map (Fig. 2D; see fig. S5F for all brain views) accounted for 34% of the variance across all 649 RSFC BWAS maps (for cortical thickness, see fig. S4D; 17% of cortical thickness map variance explained) and was nearly identical (PC_1_ *r_patterns_* = 0.97) to the SES BWAS map (Fig. 2A). The very strong overlap with SES was specific to the first PC (exposome) map for both RSFC (*r_patterns_*: PC_2_ = 0.07; PC_3_ = 0.06; fig. S11) and cortical thickness (*r_patterns_*: PC_1_ = 0.94; PC_2_ = −0.06: PC_3_ = 0.05). Some of the overlap between the exposome (Fig. 2D) and the SES (Fig. 2A) brain maps could be driven by including socioeconomic when generating the exposome map. Therefore, we recomputed the exposome map after excluding all socioeconomic variables. Doing so generated a nearly identical exposome map compared to the one that also included socioeconomic variables (*r_patterns_* = 0.99). In sum, a single, common BWAS brain pattern exists across hundreds of variables that is most reflective of a person’s socioeconomics.

To further examine the exposome map’s projection onto variable-specific BWAS maps, we correlated the exposome map with each of them (Fig. 2E; |*r_patterns_*|; fig. S4E for cortical thickness). For RSFC and cortical thickness, socioeconomic variables, on average, exhibited the strongest spatial similarity (median across all variables |*r_patterns_*|: RSFC = 0.84; cortical thickness = 0.70) to the exposome association map. Across categories, measures of screen time (median |*r_patterns_*|: RSFC = 0.67; cortical thickness = 0.42) and cognition (median |*r_patterns_*|: RSFC = 0.66; cortical thickness = 0.49) also demonstrated relatively strong similarity to the principal association map. Considering the exemplar variables with strong brain associations (SES, sleep, screen time, IQ) simultaneously in a multiple regression model (adjusting for age, sex, head motion, site; see Methods) did not alter their BWAS similarity ranking relative to the exposome association map (see Methods; fig. S12A for association strength rankings and fig. S12B-E for residualized brain maps of SES, sleep, screen time, and IQ).

The brain networks most strongly represented in the exposome BWAS pattern were early sensory (VIS, AUD), motor (SMD, SML), and the somato-cognitive action network (SCAN (*42*)) recently discovered in primary motor cortex (Fig. 2F; fig. S4F for cortical thickness). Early sensory and motor networks had significantly stronger associations with the exposome map than higher-order cognitive networks (*t* = 9.81, *P* = 9.21 × 10^−20^, one tailed independent samples t-test) for RSFC, but not for cortical thickness (fig. S4F). The frontoparietal network (*43*) (FPN; yellow), preferentially activated during complex cognition and linked to inter-individual variation in fluid IQ by task fMRI (*23*, *44*), had the least overlap with the RSFC exposome pattern (Fig. 2F).

## Comparative functional pattern analytics

To further investigate the exposome association map (RSFC), we utilized comparative functional pattern analytics, which quantifies differences and similarities across functional brain mapping modalities (Fig. 3; BWAS, task fMRI, PET receptor densities, EEG, drug trial fMRI). We derived meta-analytic task fMRI maps ((Fig. 3A, left, working memory, reasoning, cognitive control; NeuroSynth, see Methods), receptor density maps (Fig. 3A, top right; norepinephrine (brain’s primary arousal transmitter (*45*, *46*)), PET imaging; see (Supplementary Table 2 for other transmitters, all of which were less strongly correlated to the exposome map than norepinephrine), as well as maps of sleep (Fig. 3A, middle right; EEG (*47*, *48*)), stimulant use (Fig. 3A; bottom right; trial fMRI (*49*)), SES, and IQ. We quantified their spatial correlation with the exposome map. Calculation of spatial correlations across maps revealed that task fMRI contrasts of high cognitive demand were anti-correlated with the RSFC exposome map (*r_patterns_*: working memory = −0.15; reasoning = −0.17; cognitive control = −0.16) and significantly distinct from it (all *P* < 0.05, FDR corrected). In contrast, arousal variable maps were strongly correlated (all *P* < 0.05, FDR corrected) with the exposome map (*r_patterns_*: norepinephrine = 0.34; sleep = 0.95; stimulants = 0.49), as were SES (*r_patterns_* = 0.97) and IQ (*r_patterns_* = 0.78), thus revealing the IQ BWAS pattern to be mismatched with the known functional neuroanatomy of complex cognition (*44*, *50*, *51*) (Fig. 3; see fig. S13 for correlation (*r_patterns_*) between all functional brain maps with each other).

**Fig. 3.**
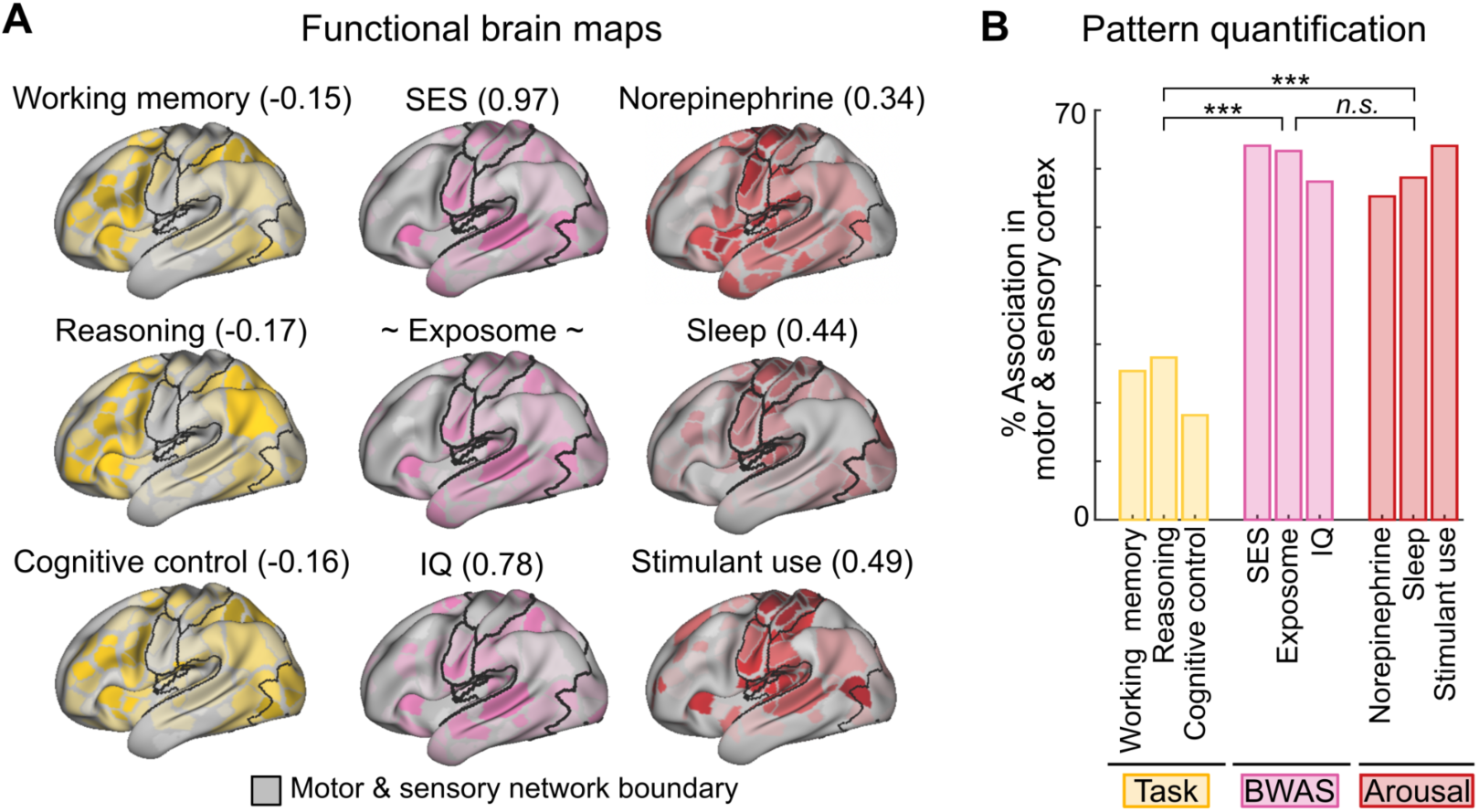
Comparative functional pattern analytics. **(A)** Comparisons of task-fMRI activations maps (gold, left) of higher-order cognitive tasks (working memory, reasoning, cognitive control; NeuroSynth), resting state functional connectivity (RSFC) BWAS maps (pink, middle; SES, exposome, IQ), and non-BWAS patterns of arousal-related effects (red, right; norepinephrine receptor density, sleep duration, stimulant use) mapped using PET (norepinephrine), EEG (sleep), and stimulant use (trial fMRI). The number next to each variable represents the correlation (*r_patterns_*) of the map to the exposome BWAS map (center). The motor and sensory networks are outlined in black. **(B)** Proportion of total association represented in motor (somatomotor dorsal, somatomotor lateral, somato-cognitive action) and sensory (visual and auditory) networks vs. all others for higher-order cognitive tasks (gold, left; working memory, reasoning, cognitive control; NeuroSynth), functional connectivity BWAS maps (pink, middle; SES, principal exposome map, IQ), and arousal-related phenotypes (red, right; norepinephrine receptor density, sleep (EEG), stimulant use). *** indicates *P* < 0.001, Bonferroni corrected. n.s.: not significant (*P* > 0.05).

To quantify how strongly each functional pattern mapped onto the brain’s primary cortex, we computed the percentage of total association strength represented in sensory and motor networks, relative to the rest of the brain (Fig. 3B). Complex cognitive fMRI task maps (working memory, reasoning, cognitive control) overlapped much less with sensory and motor (~20%) than all other (association) networks (Fig. 3B, gold, left). In contrast, the BWAS (SES, exposome, IQ; Fig. 3B, BWAS purple, middle) and arousal (norepinephrine receptor density, sleep duration, stimulant use; Fig. 3B, Arousal, red, right) maps were dominated by sensory and motor regions (~60%). A one way analysis of variance (ANOVA) comparing cognitive tasks (Fig. 3B gold, left), functional connectivity BWAS (Fig. 3B purple, middle), and arousal-related maps (Fig. 3B red, right), demonstrated that the cognition maps (Task) were significantly different from the others (BWAS, Arousal; *F*_(2,6)_ = 72.47, *P* = 6.28 × 10^−5^), while there was no statistical difference between the other sets of maps (BWAS vs. Arousal; *P* = 0.93, Tukey-Kramer multiple comparison corrected). While comparative functional pattern analytics underscored the similarity of the exposome, SES, and IQ maps with known arousal associations, it highlighted the chiasm between the accepted pre-frontal and parietal distribution of higher cognition (*52*), and the motor and sensory BWAS map of IQ (*r_patterns_* = 0.00).

## Brain with IQ associations are related to SES

It is well known from task fMRI and other methods (e.g., lesion mapping, electrophysiology, computational modeling) (*53*, *54*) that higher-order cognition (e.g., reasoning) localizes to prefrontal and parietal cortex (Fig. 3A, left). However, the BWAS pattern of IQ, which is thought to be a latent index of cognitive abilities, did not map onto prefrontal and parietal regions, but rather to sensorimotor networks (Fig. 3A, middle). One potential explanation for the BWAS IQ map not matching the known pattern of higher-order cognition is that the brain-IQ association pattern is affected by SES.

To assess the linear effects of SES on brain-wide associations with IQ, we regressed SES from IQ scores and recomputed univariate brain-wide associations with IQ (Fig. 4A,B for RSFC; fig. S14 for cortical thickness; see Methods). Adjusting for SES weakened 95% of the brain-IQ associations, for both RSFC and cortical thickness, such that ~70% of them were no longer statistically significant, while only 5% did not decrease and remained significant (*P* < 0.001, one-sided). The strongest associations between the brain and IQ systematically decreased the most when adjusting for SES (Fig. 4B; see fig. S14B for cortical thickness). Although much weaker, residualized (SES-adjusted) IQ better maps onto distributed frontoparietal regions (fig. S12E).

**Fig. 4.**
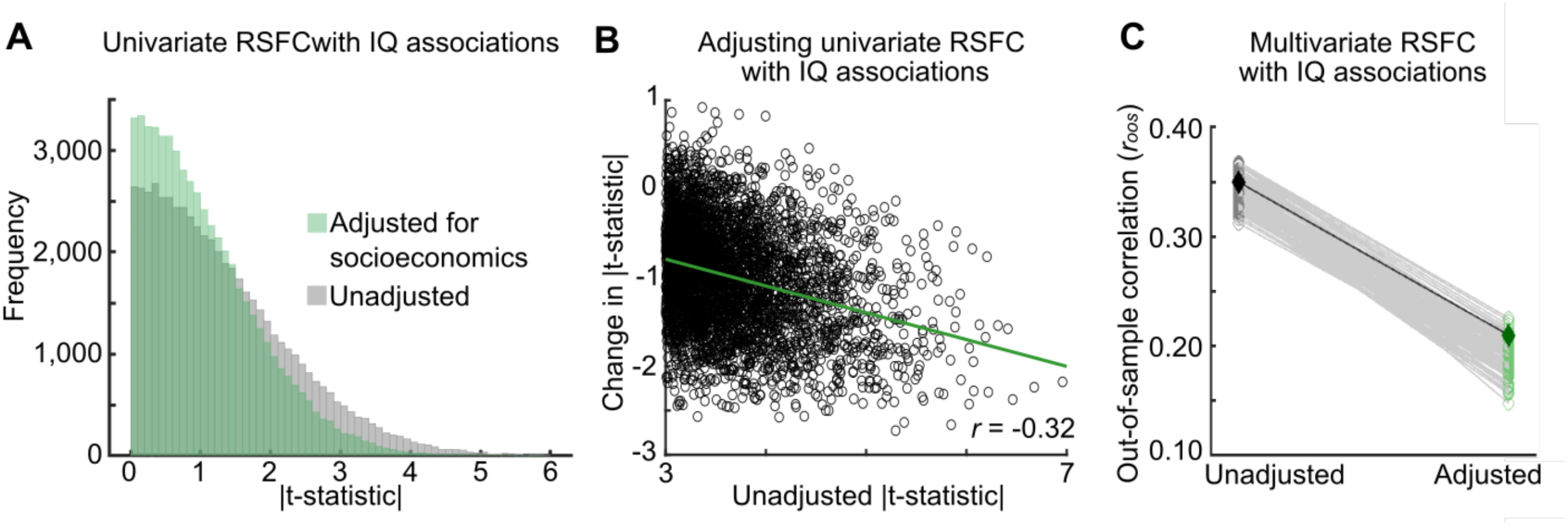
Adjusting for SES in brain-IQ associations. **(A)** Distribution of associations between resting-state functional connectivity (RSFC) edges and IQ scores (NIH Toolbox Cognition Battery, total score; see fig. S14A for cortical thickness) in univariate models unadjusted for SES (grey; social and economic domain from the Child Opportunity Index) and after adjusting for SES (green). **(B)** Univariate associations of brain function (RSFC; see fig. S14B for cortical thickness) with IQ scores after SES adjustment; (y-axis) as a function of the unadjusted association between RSFC and IQ scores (x-axis). The negative slope (*r* = −0.32) indicates that adjusting for SES reduces stronger RSFC with IQ associations the most. **(C)** Multivariate associations (out-of-sample correlations: *r_oos_*) of RSFC with IQ scores (see fig. S14C for cortical thickness) using ridge regression (see fig. S15 to S16 for replication with Connectome-Based Predictive Modeling (*55*)) unadjusted (grey; *left*) for SES and adjusted for SES (green; *right*). Models were trained and tested on 100 split-half bootstrap samples. Black diamond (left) represents the out-of-sample model fit from a predefined replication sample (see Methods), unadjusted for SES, while the dark green diamond (right) represents the out-of-sample correlation from the same predefined matched replication sample (see Methods), adjusted for SES. In all instances, the correlation between observed and predicted scores is plotted.

Multivariate associations (Fig. 4C) of the brain with IQ (unadjusted, RSFC: *r_oos_* = 0.36, in line with estimates from many previous reports in large samples (*56*); cortical thickness: *r_oos_* = 0.23) were reduced by 30-40% (adjusted, RSFC: *r_oos_* = 0.23; Fig. 4C; cortical thickness: *r_oos_* = 0.13; fig. S14C), in the same sample, after adjusting for SES when using ridge regression, and by ~50% when using Connectome-based Predictive Modeling (*55*) (CPM; fig. S15; RSFC unadjusted *r_oos_* = 0.26; adjusted *r_oos_* = 0.13; fig. S16; cortical thickness unadjusted *r_oos_* = 0.21; adjusted *r_oos_* = 0.12). As a control, multivariate associations of RSFC with SES (ridge regression) were only reduced by ~20% when adjusting for IQ (RSFC: unadjusted for IQ *r_oos_* = 0.39, adjusted for IQ *r_oos_* = 0.31; cortical thickness:: unadjusted for IQ *r_oos_* = 0.38, adjusted for IQ *r_oos_* = 0.34; fig. S17) and were similar to brain-based multivariate associations of IQ without SES adjustment (*r_oos_* = 0.32). To test for confounding of brain-IQ associations by SES with a different method, we also used confound isolation cross-validation (*57*), which tests multivariate models on subsamples without a linear association between IQ and SES. It showed SES to have stronger RSFC associations (*r_oos_* = 0.30) than IQ (*r_oos_* = 0.24; fig. S18; see Methods). Thus, using linear methods, considering SES diminishes associations between the brain and IQ scores, independent of the methodology. These results provide a potential explanation as to why the IQ BWAS pattern strongly resembles the SES BWAS pattern (i.e., confounding), but not the known patterns from task fMRI (Fig. 3a) of cortical regions most important for higher-order cognition.

## Testing the generalizability of brain-IQ associations

For brain-based multivariate models of population variability to be valid, they must generalize across cohorts (*58–61*). Hence, we tested the generalizability of multivariate brain-IQ associations across the SES spectrum. We used canonical correlation analysis (CCA), a common multivariate technique used to learn associations between multivariate brain and non-brain data, carrying forward hyperparameters from our previous work (*1*) to reduce overfitting.

We tested the generalizability of multivariate brain-IQ score association models in a so-called cross-contextual comparison framework (*6*) (Fig. 5; RSFC; for cortical thickness, see fig. S19) (*6*). Specifically, we trained brain-based models of IQ scores either on ABCD subsamples covering the full SES spectrum (Fig. 5A-E; fig. S19A-E) or subsamples restricted to either relatively high (Fig. 5F-J; social and economic domain of the Child Opportunity Index > 0.75) or low SES (Fig. 5K-O; social and economic domain of the Child Opportunity Index < −0.55) backgrounds. We subsequently tested the models using the ABCD Replication (test) sample data, matched for sample size (*n* = 569) and IQ distribution (Fig. 5A-O; fig. S19 for cortical thickness; see Methods). In both cases, the SES spectrum was truncated in the training data (Fig. 5i,n), but not the test data (Fig. 5J,O). As a baseline, the multivariate model trained on children from the full SES spectrum achieved an association of *r_oos_* = 0.27 in the test subsample (*P* < 0.001; Fig. 5a; fig. S19 for cortical thickness; figs. S20 to S21 for CPM). However, the model trained on only higher SES children SES failed to generalize out-of-sample (IQ-matched), with an *r_oos_* = 0.01 (*P* = 0.45; Fig. 5F). Conversely, the model trained on only lower SES children (IQ-matched) achieved a strong association of *r_oos_* = 0.34 in the test subsample (*P* < 0.001; Fig. 5K). Repeating these analyses with cortical thickness generated the same results (fig. S19). The failure of brain-IQ associations to generalize when models were trained on individuals from high, but not low, SES backgrounds, suggests a potential non-linear SES dependence of brain-IQ associations.

**Fig. 5.**
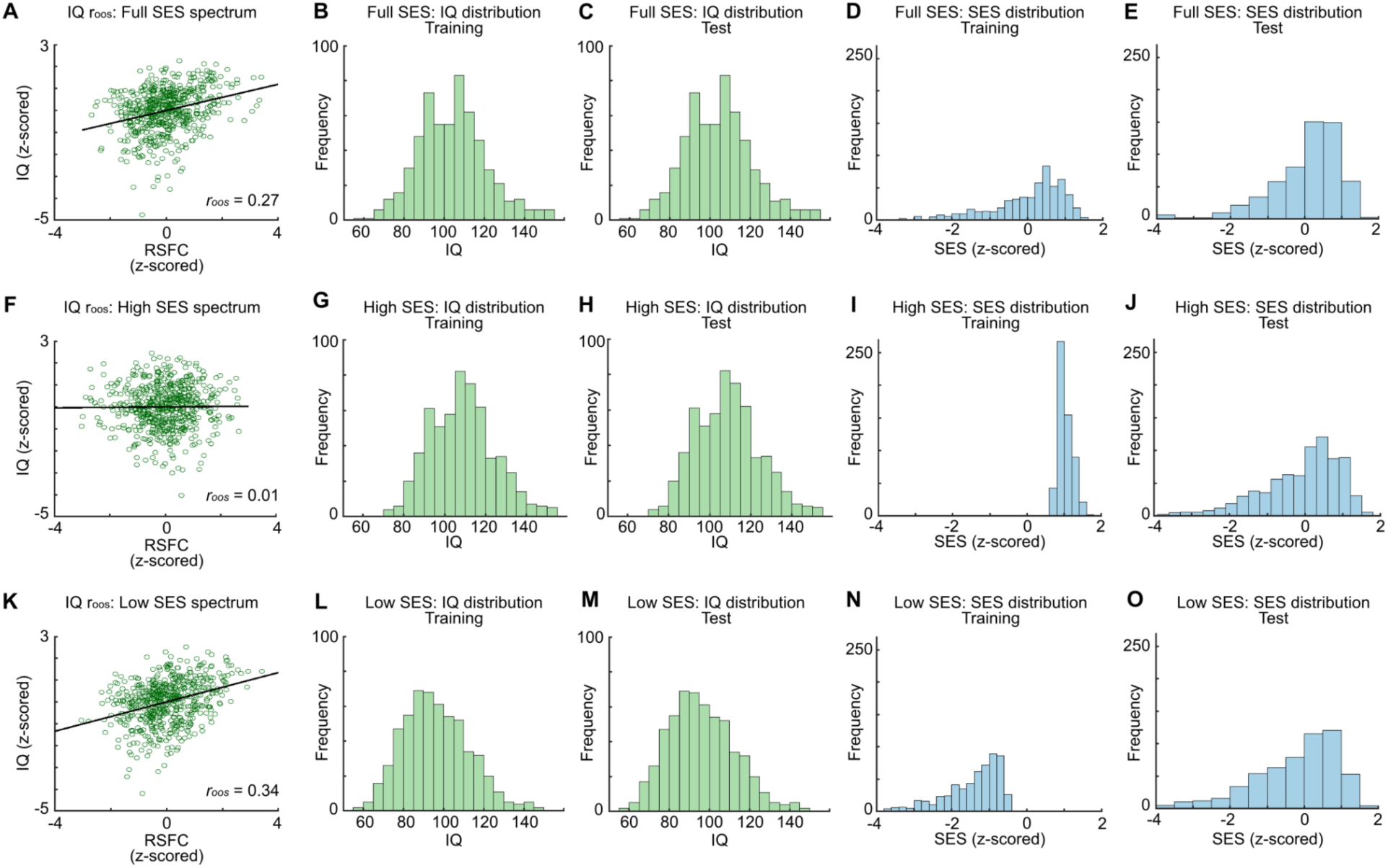
Non-generalizability of multivariate brain-IQ score associations. **(A)** Out-of-sample IQ (NIH Toolbox Cognition Battery, all subscales) associations from resting-state functional connectivity (RSFC, 333 parcels; see fig. S19 for cortical thickness), using canonical correlation analysis (CCA), in the Adolescent Brain Cognitive Development (ABCD) Study. The y-axis shows the first canonical variates for IQ in the test data, scaled by the training weights. The x-axis shows the same but for RSFC. The correlation between the first canonical variates for IQ and RSFC is out-of-sample relative to the training data (*r_oos_*). The training and test sets were subsampled from the pre-defined ABCD Discovery (*n* = 2,316) and Replication (*n* = 2,263) samples, respectively, thus covering the full ABCD SES spectrum (social and economic domain of the Child Opportunity Index) and matched for size (*n* = 569). Subsample multivariate associations (*r_oos_* = 0.27; *n* = 569) were lower than in the complete sample (*r_oos_* = 0.35; *n* = 2,316) due to the known scaling of effect size with *n*. **(B)** IQ distribution in the training sample (full SES) **(C)** was matched to the test sample (full SES). This led to similar SES distributions in the **(D)** training and **(E)** test samples. **(F)** Out-of-sample IQ score with RSFC association, exactly as in **(A)**, but for a size-matched (*n* = 569) training subsample drawn from only the high SES spectrum (z > 0.75). Restricting the training sample to high SES, while keeping the test sample as in **(A)**, (full SES) reduced multivariate association strength to *r_oos_* = 0.01 (*P* = 0.45). **(G)** IQ distribution in the training sample (high SES) was matched to **(H)** the test sample (full SES). This restricted the **(I)** training SES distribution to high, **(J)** but not for the test sample. **(K)** Out-of-sample IQ score associated with RSFC association, exactly as in a, but for a size-matched (*n* = 569) training subsample drawn from only the low SES spectrum (z < −0.55). Restricting the training sample to low SES, while keeping the test sample as in a (full SES) increased multivariate association strength to *r_oos_* = 0.34. **(L)** IQ distribution in the training sample (low SES) was matched to **(M)** the test sample (full SES). This restricted the **(N)** training SES distribution to low, **(O)** but not for the test sample.

## Shortcut learning in brain-IQ associations

To determine why generalizability plummeted as subsamples became more restricted to higher SES children, we implemented a more fine-grained cross-contextual analysis (*6*). We continuously varied the SES composition of the training sample, starting with children from only high SES backgrounds, and gradually moved to include the full SES spectrum. These models were then tested on IQ-matched replication samples (Fig. 6A), while tracking the correlation (*r_vars_*) between SES and IQ. As training samples progressed from most SES-restricted (high only) to least restricted, multivariate subsample RSFC associations increased from *r_oos_* = 0.03 to *r_oos_* = 0.23 (Fig. 6A; fig. S22A for cortical thickness; *r_oos_* = 0.02 to *r_oos_* = 0.16). Paralleling this trajectory, the correlation between SES and IQ in the training subsamples increased from *r_vars_* = 0.08 in high-SES subsamples to *r_vars_* = 0.31 when subsampling the full cohort (Fig. 6B, fig. S22B for cortical thickness). RSFC and cortical thickness-based models of IQ only succeeded when the correlation between SES and IQ was at least *r_vars_* > 0.10. Despite family income and parental education (family level socioeconomic variables) also having strong correlations with IQ scores (*r_vars_* = 0.34 for both), varying the training sample by them substantially reduced multivariate associations, but less than for SES (fig. S23). This indicates that neighborhood level socioeconomic variables have an especially strong effect on brain-IQ associations.

**Fig. 6.**
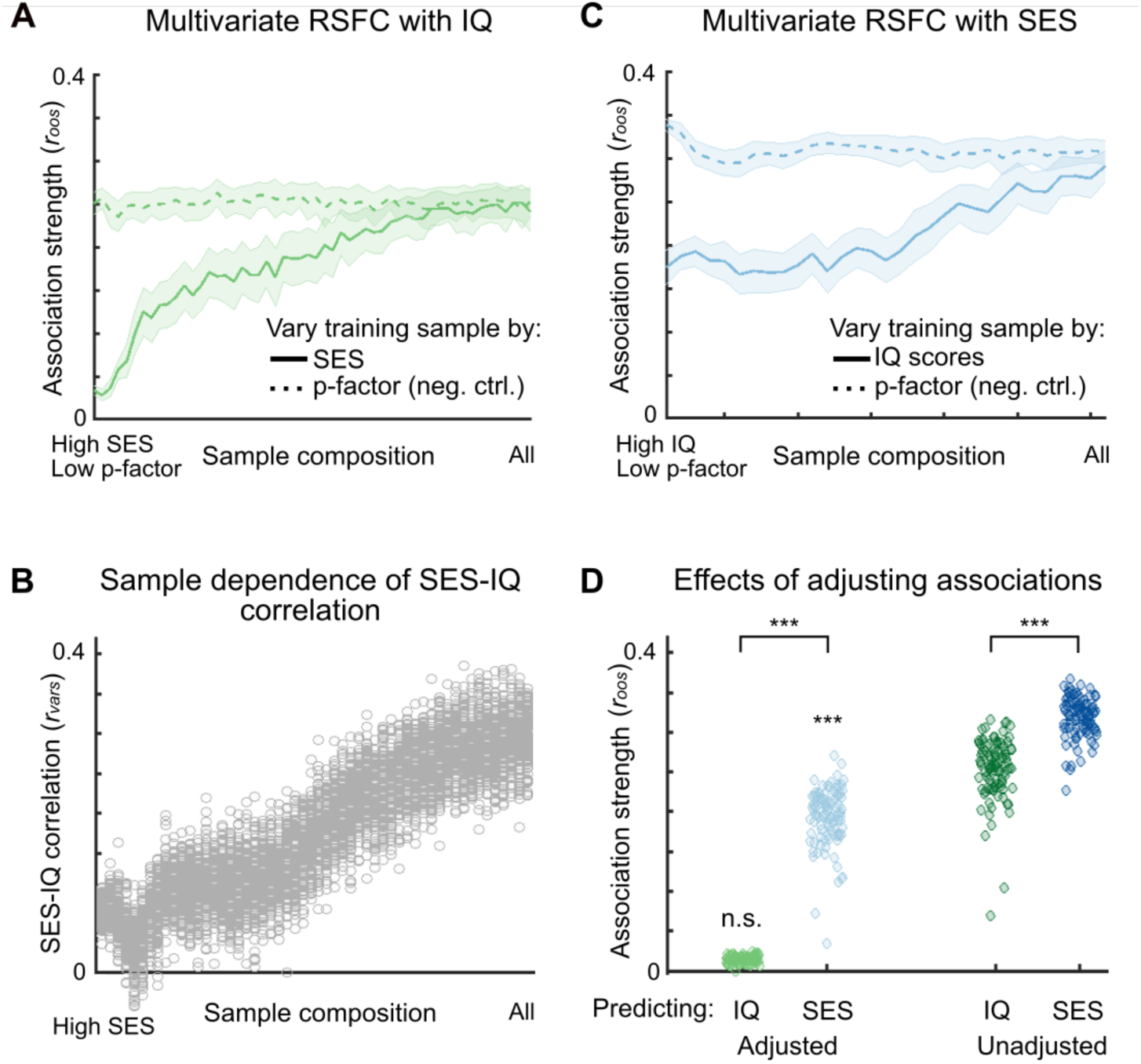
Cross-contextual analyses testing for shortcut learning in multivariate brain associations with IQ and SES. **(A)** IQ (NIH Toolbox Cognition Battery, total score) was associated with resting-state functional connectivity (RSFC, 333 parcels), using canonical correlation analysis (CCA), in the Adolescent Brain Cognitive Development (ABCD) Study (as in Fig. 5), with training data of varying SES compositions. The pre-defined ABCD Discovery (*n* = 2,316) data were repeatedly subsampled (*n* = 569) to range from subsamples from the full dataset (x-axis, right) to restricted (left; high SES), based on SES (social and economic domain of the Child Opportunity Index; light blue), and psychopathology (p-factor; dark green; negative control). Multivariate associations, measured as out-of-sample correlation (*r_oos_*) using the ABCD Replication sample (*n* = 2,263), are shown on the y-axis; line shading indicates one standard deviation (SD) around the mean *r_oos_* across 100 bootstrapped subsamples. **(B)** For the brain-based multivariate associations of IQ, varying by SES in **A**, the underlying correlations between SES and IQ (*r_vars_*, y-axis; light blue), are shown as a function of SES composition of subsamples. **(C)** SES was associated with RSFC (333 parcels), using CCA, in the ABCD (Baseline), with training data of varying SES compositions. The Discovery (*n* = 2,316) data were repeatedly subsampled (*n* = 569, 31 bins, 100 bootstraps per bin) to range from distributions from the full sample (x-axis, right) to restricted (left), based on IQ (light green; high IQ on left). Varying the training sample by psychopathology (p-factor; dark green), again served as a negative control. Multivariate associations (*r_oos_*) are shown on the y-axis; line shading indicates one SD around the mean *r_oos_* across 100 bootstrapped samples. **(D)** RSFC multivariate out-of-sample association of IQ fell to *r_oos_* = 0.01 (light green dots; IQ adjusted; *P* = 0.43, not significantly different from zero, mean across 100 bootstrapped samples; for cortical thickness, see fig. S22) in subsamples in which there was a small correlation between SES and IQ (*r_vars_* < 0.10; Fig. 6B). In subsamples with a stronger correlation between SES and IQ consistent with the ABCD sample (*r_vars_* ~ 0.30), multivariate associations of IQ averaged *r_oos_* = 0.24 (*P* < 0.001; dark green dots; IQ unadjusted). Brain-based (RSFC) multivariate associations of SES remained robust at *r_oos_* = 0.19 (light blue dots; SES adjusted; *P* = 2.54 × 10^−23^) in subsamples in which there was a small correlation between SES and IQ (*r_vars_* < 0.10). In subsamples with a stronger correlation between SES and IQ consistent with the ABCD sample (*r_vars_* ~ 0.30), RSFC out-of-sample associations of SES averaged *r_oos_* = 0.32 (dark blue dots; SES unadjusted).

To test whether brain-based associations of SES were more robust to confounds than those for IQ, we evaluated multivariate models (Fig. 6C for RSFC; fig. S22C for cortical thickness) by training them on ABCD subsamples ranging from high IQ (NIH Toolbox Cognition Battery, total score > 110) only (Fig. 6C, left, lighter green line; fig. S22C for cortical thickness) to the full IQ spectrum in ABCD (Fig. 6C, right, lighter green line; fig. S22C for cortical thickness). Across all sample compositions (Fig. 6C; x-axis), generalizability remained greater for brain-based models of SES (all *r_oos_* > 0.20; Fig. 6c, lighter green line) than IQ (Fig. 6A, lighter blue line). Even when the correlation between SES and IQ had been reduced to *r_vars_* < 0.10 (Fig. 6B; fig. S22B for cortical thickness), association strength remained high (Fig. 6C,D all *r_oos_* > 0.20; fig. S22C,D all *r_oos_* > 0.15). Moreover, SES associations remained strong when varying the training sample by genetic ancestry (fig. S24, grey line; see fig. S25 for each genetic principal component; see Methods). Thus, unlike brain-IQ associations, which did not generalize (*r_oos_* = 0.01, *P* = 0.43; Fig. 6D) without a correlation (*r_vars_*) with SES, multivariate brain-SES associations remained strong (*r_oos_* = 0.19, *P* = 2.54 × 10^−23^; Fig. 6D; fig. S22 for cortical thickness).

As a negative control, we also varied the training sample composition by psychopathology (p-factor; Fig. 6A; darker green line), when predicting IQ scores (Fig. 6A) and SES (Fig. 6C) from RSFC data. Stratification of the training samples by psychopathology (starting with a low p-factor) had no effect on IQ and SES association generalizability (Fig. 6C, dark green lines; fig. S26 for IQ score non-generalizability in the Human Connectome Project (HCP)). As another negative control, we also showed that multivariate brain-based IQ and SES associations generalized across sexes, by training on female children and testing on males (*r_oos_* = 0.31) and vice versa (*r_oos_* = 0.29; fig. S27; for SES generalizability across sex, see fig. S28).

IQ models trained on children from high SES in the ABCD sample (left-most side of Fig. 6A,C, x-axis) also exhibited low generalizability to the Human Connectome Project (HCP; *n* = 877, aged 22-35 years) dataset (fig. S26; *r_oos_* = 0.03). As ABCD training models increasingly included children from lower SES backgrounds, generalizability to the HCP sample similarly improved (fig. S26; *r_oos_* ~0.20).

Thus, the generalizability of multivariate brain-based (RSFC and cortical thickness) association models with IQ scores varies with SES, independent of analytic approach (Fig. 5,6, fig. S19 to S22, fig. S24 to S26). In samples restricted to children with high SES, in which only a negligible or weak correlation between SES and IQ scores existed (*r_vars_* < 0.10), out-of-sample brain-IQ score associations were small (*r_oos_* < 0.10; Fig. 5F; Fig. 6A, solid green line, left side of plot; Fig. 6D). Multivariate associations between the brain and IQ were only observable when a bivariate correlation between IQ and SES exists.

## The socioeconomic brain pattern

Brain associations with SES were found in sensory and motor regions, in a pattern most similar to that of norepinephrine receptors, sleep correlates tracked with EEG, and the effects of high-dose methylphenidate (Fig. 3). Notably, significant overlap between the SES map and neurotransmitter maps was specific to norepinephrine (*P* = 0.002), as no other neurotransmitter maps were significantly similar to SES (all *P’s* > 0.05; Supplementary Table 2).

Like SES, sleep, screen time, and IQ BWAS maps were patterned onto early sensory and motor networks. This network pattern, present in RSFC data, does not correspond to brain regions active during complex cognitive operations (frontoparietal network (*43*), Figs. 2,3), such as, manipulating information held in working memory (Fig. 3). Rather, this network pattern exhibits the greatest day-to-day variability (*9*) within an individual and changes with drastic motor interventions (*62*), sleep deprivation (*19*), caffeine intake (*17*, *63*), length of day (*64*) (seasonal effects), stimulant use, and high norepinephrine receptor density (Fig. 3; fig. S29), altogether demonstrating its environmental sensitivity. Given that an individual’s environmental exposures have powerful effects on the brain’s neuroendocrine system (*65*), the observed principal BWAS map may be representing a biological pathway through which socioeconomics is becoming biologically embedded via differences in screen use (in particular social media (*66*)), sleep quality (*67*), and chronic physiological stress (*68*). To give credence to this abductive argument, we compiled a composite of stress in the ABCD Study using measures established previously (*69*) related to abuse, household challenges, and neglect (see Methods). The correlation between the stress RSFC brain map pattern and the SES brain pattern was *r_patterns_* = 0.50 (P < 0.001). Thus, the SES brain pattern may reflect the direct effects of childhood sleep deprivation and stress, as well as responsive adaptations. Although the developing brain adjusts to its environment, socioeconomic brain patterns similar to those associated with detrimental factors, such as insufficient sleep (*70*) and chronic stress (*68*), are unlikely to be adaptive longer term.

Childhood poverty is correlated with poorer physical and mental health outcomes, lower lifetime earnings, and decreased life expectancy (*71–74*). Gaining deeper insight into modifiable variables through which SES affects brain development (*75*) is critically important. In addition to sleep and screen time, there are numerous other pathways through which socioeconomics likely impact brain development, as well as cognitive and psychiatric outcomes (*76*, *77*). Neighborhood contributions capture individual differences in many potentially influential factors, such as schools (*78*), threat/crime (*79*), and access to healthcare (*80*). Other pathways include nutrition (*81*), physiological stressors (*82*, *83*), inflammation (*84*), and pollution (e.g., lead) (*85*).

## Reification of IQ into the brain

One goal of inquiries into the neurobiological basis of IQ has been to attribute unique and generalizable predictive power to brain features that explain individual differences in IQ scores. Generalizability of brain-based IQ models required including children with lower SES (Fig. 5, 6A, fig. S19, fig. S22A). Machine learning techniques, though powerful and more sensitive than univariate approaches, can detect associations outside of the target variable (here, IQ) that lack generalizability, replicability, or practicality (*86*). In medical applications, machine learning has generated invalid findings by learning background information of images related to lungs with Covid-19, breast cancer, and cell types, instead of the target variables (*87*), a phenomenon known as shortcut learning (*88*). Similarly, here, brain-based multivariate models of IQ were dependent on the sample’s socioeconomic background. This suggests that similar to previous work in medical imaging, multivariate brain-based models are learning socioeconomic background as a shortcut, rather than the intended target of IQ.

BWAS of IQ highlight the importance of taking the effects of childhood socioeconomics into consideration. The generalizability of brain-IQ associations was dependent on socioeconomics, similar to the Scarr-Rowe effect in genetics where the heritability of IQ is dependent on an individual’s SES (*89*). In BWAS, brain with IQ associations did not generalize when models were trained on only higher SES individuals, despite a typical distribution of IQ scores. This finding along with previous studies (*90*) calls into question the degree to which IQ scores measure a stable, essentialized trait. Amongst immigrants, IQ scores within individuals increased with time lived in the US (*91*). Moreover, IQ scores are higher for individuals adopted out of institutional care compared to those who were not (*92*). IQs also increased over much of the 20^th^ century (Flynn effect) (*93*). However, population estimates of IQ in many developed countries peaked in the late 20^th^ century, subsequently reversed (*94*, *95*), and can be explained by environmental causes (*95*).

## Environment matters for a child’s brain

The socioeconomic opportunities provided by a child’s environment are associated with brain function and structure more than any other variable examined, suggesting it captures the principal vector of the exposome, along which lived experience manifests in brain organization. Despite having the strongest effect sizes, SES still only explained a subset of brain differences across children (at maximum, 16% with multivariate approaches). Population-level effects cannot predict future outcomes of an individual child. Socioeconomic opportunity is not destiny.

Leading candidates for cost-effectively and rapidly bolstering brain function and structure may be lifestyle interventions related to sleep and chronic stress. Within-person, longitudinal, clinical trials are needed to test for robust developmental benefits of sleep and physiological stress interventions. The association patterns between IQ and primary motor and sensory regions, indicate that prior brain-IQ correlations may have been driven by physiological stressors related to lower SES (Fig. 3). Claims of IQ tests measuring essentialized intelligence are not neurobiologically grounded. While correlational, the BWAS maps are nonetheless highlighting the great power of environmental stressors and deprivation, further adding to the worry voiced by Charles Darwin, that “if the misery of the poor not be caused by the laws of nature, but by our institutions, great be our sin (*96*).”

## Acknowledgments

We thank Kathryn Humphries for thoughtful comments on our manuscript.

## ABCD

Data used in the preparation of this article were obtained from the Adolescent Brain Cognitive Development (ABCD) Study (https://abcdstudy.org), held in the NIMH Data Archive (NDA). This is a multisite, longitudinal study designed to recruit more than 10,000 children age 9-10 and follow them over 10 years into early adulthood. The ABCD Study is supported by the National Institutes of Health and additional federal partners under award numbers U01DA041022, U01DA041028, U01DA041048, U01DA041089, U01DA041106, U01DA041117, U01DA041120, U01DA041134, U01DA041148, U01DA041156, U01DA041174, U24DA041123, U24DA041147, U01DA041093, and U01DA041025. A full list of supporters is available at https://abcdstudy.org/federal-partners.html. A listing of participating sites and a complete listing of the study investigators can be found at https://abcdstudy.org/scientists/workgroups/. ABCD consortium investigators designed and implemented the study and/or provided data but did not necessarily participate in analysis or writing of this report. This manuscript reflects the views of the authors and may not reflect the opinions or views of the NIH or ABCD consortium investigators.

## Human Connectome Project Study

Data were provided, in part, by the Human Connectome Project, WU-Minn Consortium (U54 MH091657) funded by the 16 NIH Institutes and Centers that support the NIH Blueprint for Neuroscience Research; and by the McDonnell Center for Systems Neuroscience at Washington University.

## UK Biobank Study

This research has been conducted, in part, using data from UK Biobank (www.ukbiobank.ac.uk). UK Biobank is generously supported by its founding funders the Wellcome Trust and UK Medical Research Council, as well as the Department of Health, Scottish Government, the Northwest Regional Development Agency, British Heart Foundation and Cancer Research UK.

## Daenerys NCCR

This work used the storage and computational resources provided by the Daenerys Neuroimaging Community Computing Resource (NCCR). The Daenerys NCCR is supported by the McDonnell Center for Systems Neuroscience at Washington University, the Intellectual and Developmental Disabilities Research Center (IDDRC; P50 HD103525) at Washington University School of Medicine and the Institute of Clinical and Translational Sciences (ICTS; UL1 TR002345) at Washington University School of Medicine.

This manuscript is the result of funding in whole or in part by the National Institutes of Health (NIH). It is subject to the NIH Public Access Policy. Through acceptance of this federal funding, NIH has been given a right to make this manuscript publicly available in PubMed Central upon the Official Date of Publication, as defined by NIH.

The views expressed are those of the authors and do not necessarily represent the official views of the National Institutes of Health.

## Author contributions

Conceptualized study design and methodology: SM, BTC, NUFD

Data curation, analysis, and code: SM, MRD, NK, RJC, ACM, JM, ANV, VS, SEP, AJG, TJH, SMM

Writing of original draft: SM, MRD, DAF, BTC, NUFD

All authors reviewed, provided comments, and edited the final manuscript

## Data and materials availability

Participant level data from all datasets (ABCD, HCP, UK Biobank) are openly available pursuant to individual, consortium-level data access rules. The ABCD data repository grows and changes over time (https://nda.nih.gov/abcd). The ABCD data used in this report came from ABCD collection 3165 and the Annual Releases 3.0,4.0, and 5.1; DOI 10.15154/1503209.

Data were provided, in part, by the Human Connectome Project, WU-Minn Consortium (Principal Investigators: David Van Essen and Kamil Ugurbil; 1U54MH091657) funded by the 16 NIH Institutes and Centers that support the NIH Blueprint for Neuroscience Research; and by the McDonnell Center for Systems Neuroscience at Washington University. Some data used in the present study are available for download from the Human Connectome Project (www.humanconnectome.org). Users must agree to data use terms for the HCP before being allowed access to the data and ConnectomeDB, details are provided at https://www.humanconnectome.org/study/hcp-young-adult/data-use-terms.

The UK Biobank is a large-scale biomedical database and research resource containing genetic, lifestyle and health information from half a million UK participants (www.ukbiobank.ac.uk). UK Biobank’s database, which includes blood samples, heart and brain scans and genetic data of the 500,000 volunteer participants, is globally accessible to approved researchers who are undertaking health-related research that is in the public interest.

Analysis code specific to this study is available here: https://gitlab.com/DosenbachGreene/ses_nature

Code for processing ABCD and UKB data can be found here: https://github.com/DCAN-Labs/abcd-hcp-pipeline

