## Supplemental Information for "Patterns of brain-wide associations reflect socioeconomics"

#### Supplementary Materials

#### Materials and Methods

This study received approval from the institutional review board from each of the ABCD sites. Ethical regulations were followed during data collection and analysis. Parents and/or caregivers provided written informed consent, and children gave written assent.

Datasets

*Adolescent Brain Cognitive Development (ABCD) dataset*

Non-imaging variables and neuroimaging (resting-state functional connectivity, cortical thickness) data were from the ABCD (5.1 release; Fast Track Release 3165 collection) dataset (*n* = 4,579; see ref[1] for details on participant information), the Human Connectome Project HCP (HCP; 1200 Subjects Data Release) dataset (see below), and the UK Biobank dataset (see below). The ABCD Study obtained centralized institutional review board (IRB) approval from the University of California, San Diego. Each of the 21 sites also obtained local IRB approval. Ethical regulations were followed during data collection and analysis. Parents or caregivers provided written informed consent, and children gave written assent.

Here, we used ABCD Study data from the Baseline (9-10 year old) sample in all primary analyses. Univariate associations between RSFC and non-imaging variables were replicated using data from the Year 2 wave of ABCD Study (see below). For primary analyses leveraging the Baseline ABCD sample, cortical thickness and resting-state functional connectivity data from *n* = 10,259 participants were obtained through the ABCD fast track portal. To obtain the final sample size, children from the full behavioral sample (*n* = 11,878) were first divided into a Discovery (*n* = 5,939) and Replication (*n* = 5,939) set, which were previously matched(*40*) across nine variables using the ABCD 3.0 data release: site location, age, sex, ethnicity, grade, highest level of parental education, handedness, combined family income, and exposure to anesthesia. Family members (e.g., sibling pairs, twins, and triplets) were kept together in the same set and the two sets were matched to include equal numbers of single participant and family members. Please refer to fig. S30 for a consort diagram of the data sets.

For supplementary analyses using Year 2 ABCD Study data (aged 11-12 years), RSFC data from *n* = 5,386 participants were obtained through the ABCD fast track portal. Phenotypic assessments collected in Year 2 along with the linked external data from the Baseline sample were used for replication of effect sizes (384 variables in total).

Head motion can systematically bias developmental studies (*97*, *98*), as well as those relating RSFC to behavior (*97*). However, these systematic biases can be addressed through rigorous head motion correction (*99*). Therefore, we used strict inclusion criteria with regard to head motion in the current study. Specifically, inclusion criteria for the current project (see (*100*) for broader ABCD inclusion criteria) consisted of at least 365 frames (5 minutes) of low-motion (FD < 0.08) RSFC data in the ABCD dataset (8 minutes of FD < 0.20 for HCP). Based on these criteria, our final Baseline ABCD dataset consisted of data from a total of *n* = 4,579 youth across the Discovery (Arm1; *n* = 2,316) and Replication (Arm2; *n* = 2,263) sets. The final 2 year follow up ABCD dataset consisted of RSFC data from 2,363 individuals. The final HCP dataset consisted of $n = 877$ individuals aged 22-35 years. The final UK Biobank dataset consisted of $n = 32,572$ individuals, aged 40-69 years.

*Human Connectome Project (HCP) dataset*
We used data from $n = 877$ individuals from the HCP 1,200 Subject Data Release (aged 22-35 years). All HCP participants provided informed consent. A custom Siemens SKYRA 3.0T MRI scanner and a custom 32-channel Head Matrix Coil were used to obtain high-resolution T1-weighted (MP-RAGE, $TR = 2.4$ s, 0.7 $mm^3$ voxels) and BOLD contrast sensitive (gradient-echo EPI, multiband factor 8, $TR = 0.72$ s, 2 $mm^3$ voxels) images from each participant. The HCP used sequences with left-to-right (LR) and right-to-left (RL) phase encoding, with a single RL and LR run on each day for two consecutive days for a total of four runs[68]. MRI data were preprocessed as previously described[62]. All HCP data are available at https://db.humanconnectome.org/.

Similar to the ABCD data, we extracted the timeseries from a total of 333 cortical regions of interest, and subsequently correlated and Fisher *z*-transformed them. We tested generalizability of multivariate models trained on ABCD data using vectorized RSFC data, as well as data from all subscales of the NIH Toolbox Cognition Battery. For each iteration across all sampling bins, RSFC edges were projected through the principal component space of the ABCD training data.

*UK Biobank (UKB) dataset*
We used pre-processed resting-state data from $N = 37,000$ individuals from the January 2020 UK Biobank release (*101*). For a complete description of study flow and imaging protocols, see (*8*). Similar to the ABCD data, we extracted the timeseries from a total of 333 cortical regions of interest, and subsequently correlated and Fisher *z*-transformed them. The UK Biobank collects measures of fluid intelligence and income, which we used to correlate with RSFC, mimicking the ABCD.

#### MRI acquisition

Imaging for each ABCD youth was performed across 21 sites within the United States, harmonized across Siemens Prisma, Philips, and GE 3T scanners. Details on image acquisition can be found in(*100*). Twenty minutes of eyes-open (passive crosshair viewing) resting state data were presented to ensure at least 5 minutes of low-motion data. All resting state scans were acquired using a gradient-echo EPI sequence (TR = 800 ms, TE = 30 ms, flip angle = 90°, voxel size = 2.4 $mm^3$, 60 slices). Head motion was monitored online using Framewise Integrated Real-time MRI Monitor (FIRMM) software at Siemens sites (*102*).

#### Neuroimaging data processing overview

We outline the data processing steps here. For further details on the full processing stream see ref(*40*). The ABCD-HCP pipeline comprises six stages: (1) PreFreeSurfer, which normalizes anatomical data; (2) FreeSurfer, which constructs cortical surfaces from the normalized anatomical data; (3) PostFreeSurfer, which converts outputs from FreeSurfer to CIFTIs and transforms the volumes to a standard volume space using ANTs nonlinear registration; (4) "Vol" stage, which performs the atlas transformation, mean field distortion correction, and resampling to 2-mm isotropic voxels in a single step using FSL's applywarp tool; (5) "Surf" stage, which projects the volumetric functional data onto the surface; and (6) "DCANBOLDproc", which performs functional connectivity processing.

DCANBOLDproc includes a respiratory filter to improve FD estimates calculated in the “vol” stage. Second, temporal masks were created to flag motion-contaminated frames using the improved FD estimates(*98*). Frames with FD > 0.30 mm were flagged as motion-contaminated. After computing the temporal masks for high motion frame censoring, the data were processed with the following steps: (i) demeaning and detrending, (ii) interpolation across censored frames using least squares spectral estimation of the values at censored frames(*99*) so that continuous data can be (iii) denoised via a GLM including: whole brain, ventricular, and white matter signals, as well as their derivatives. Denoised data are then passed through (iv) a band-pass filter (0.008 Hz < f < 0.10 Hz) without re-introducing nuisance signals (*103*) or contaminating frames near high motion frames (*104*).

For each participant, RSFC matrices were created by correlating the average time series of BOLD activity within each of 333 cortical ROIs/parcels after discarding frames with an FD > 0.08. For each participant, cortical thickness was extracted from 59,412 cortical vertices.

For additional details related to MRI acquisition and processing for ABCD and HCP data, see (*1*). In each dataset (HCP, ABCD), we extracted the timeseries from a total of 333 cortical ROIs, correlated and Fisher *z*-transformed them.

<u>Phenotypic assessment</u>

In ABCD, we assessed 649 variables, spanning 12 categories: socioeconomics, screen time, cognition, demographics, physical health, mental health, culture/environment, social adjustment, substance use, parenting, personality, and medical history. A complete list of variables and their effect size rank can be found in Supplementary Table 1. The measures included in each domain are briefly summarized below.

*Socioeconomic*. We used measures from the census tract produced by the Linked External Data (LED) Environmental workgroup (*105*), and family income to estimate socioeconomics. Family income includes wages, benefits, and child support payments. The LED incorporates the use of geospatial location data to assess individual, neighborhood, and state level data based on the child’s residential (census tract) location. See reference (*105*) for a detailed description of the LED. Notable linked datasets include the Area Deprivation Index, the Child Opportunity Index, and the Opportunity Atlas.

*Screen Time.* Caregiver- and youth-reported screen time metrics were included. The questionnaires ask about time spent on screens for various types of media, separated for weekdays and weekend days, as well as the frequency of certain screen media activities (e.g., mature-rated video games).

*Cognition.* All available age-corrected summary scores from the NIH Toolbox Cognition Battery were included, as were summary scores for the Little Man Task and Rey Auditory Verbal Learning Task. Matrix Reasoning and the Cash Choice Task measures were also included.

*Demographics.* Caregiver-reported demographic variables, such as education, employment, marital status, among others, were included. Family income was also obtained and was categorized as a socioeconomic variable.

*Culture/Environment.* Cultural and environmental variables included the neighborhood safety and environment, and youth prosocial behavior; and school risk and protective factors; and neighborhood environment measures as assessed on the residential history derived scores, such as levels of lead, ozone, and other toxins.

*Physical Health.* Physical health measures consisted of caregiver-reported sleep disturbances, youth- and caregiver-reported pubertal development, traumatic brain injury information, sports and activities involvement, and anthropometrics (i.e., birth weight, height, weight, waist circumference).

*Mental Health.* Mental health consists of items reflecting family history and caregiver symptoms and behaviors, as well as child symptoms and behaviors. Measures included the CBCL, Brief Problem Monitor-Teacher Report, Prodromal Psychosis Scale, the General Behavior Inventory-Mania, Resilience, and the Bis/Bas System Scales. The KSADS-5 was also administered.

*Social Adjustment.* Social adjustment consists of questionnaires related to peer influence, resilience, socialization, discrimination, and social influence and understanding.

*Substance Use.* Youth self-reported substance familiarity and use patterns, as well as intention to use, subjective response to various substances, and peer use patterns were assessed. Further, objective measures of use, including breathalyzer, nicalert, hair tests, and other toxicology tests are available for all or a subset of participants. Finally, measures of parental rules about substance use and community risk and protective factors were also assessed.

*Personality*. Personality was assessed by a caregiver report using the Urgency-Premeditation-Perseverance-Sensation Seeking-Positive Urgency (UPPS-P) scale.

*Parenting.* Youth- and caregiver-reported family environment scale, youth-reported parental monitoring, children's report of parental behavior inventory.

*Medical history.* Medical history measures consisted of variables related to medical history (e.g., broken bones, ER visits), medication inventory, and a developmental history questionnaire, which assesses prenatal, pregnancy, and birth events and exposures, and pubertal hormones (i.e., testosterone, estradiol, and DHEA).

Race and genetic ancestry

The National Academies of Science, Engineering, and Medicine (NASEM) issued a report providing recommendations on using race, ethnicity, ancestry, and other descriptors of population stratification (*106*). The NASEM report asserts that race does not have any biological basis and emphasizes that researchers not use continental labels (that is, African, European, Asian, Native American, and Oceanian ancestry) or socially constructed racial and ethnic categories in genetics studies (Asian, Black, White, etc.). Similarly, for neuroimaging studies, researchers should not blindly "control for race" in brain-based association studies with non-brain variables, especially when race and ethnicity are patterned onto said variables (*107–109*). We acknowledge the complicated historical and present social and economic backdrop in which race, ethnicity, and SES co-occur.

The current study follows these recommendations, and therefore does not evaluate race and ethnicity as target variables in univariate and multivariate brain association models (*110*, *111*) Genetic ancestry is defined within the context of genetic admixture of specific, idiosyncratic reference populations (in this case, ABCD). Ancestral inference methods (principal component analysis in ABCD), while more robust than self-reported race, can still be biased by the sampling process and model parameters (*112*). Therefore, we leveraged the UK Biobank to ameliorate concerns regarding race, since the UK Biobank is relatively more homogenous than the ABCD Study with respect to race (95% White British, White Irish, or other White background). With regard to genetic ancestry in the ABCD Study, we varied the training samples in brain-based (RSFC) multivariate models of SES by genetic ancestry to establish the robustness of brain with SES models to potential differences in genetic ancestry. Neither race nor genetic ancestry accounted for the observed socioeconomic effects or brain-wide association patterns (fig. S7, fig. S24).

Data Analysis

All models and data shown in the main text were tested for replication in the independent Baseline Replication (Arm2) ABCD sample (*40*).

*Manhattan plots*

Univariate analyses: We correlated each of the 649 non-brain variables with each region-of-interest pair (edge) in the resting-state functional connectivity (RSFC) data, separately. This procedure resulted in 55,278 associations between RSFC and each non-brain variable. The effect sizes of all associations are plotted in Fig. 1, sorted by the 99$^{th}$ percentile (confidence interval; 99% CI) across all brain features within each of the 649 variables. Rankings were highly similar when using the maximum or 95% CI $r_{BWAS}$ as the threshold instead of the 99% CI $r_{BWAS}$ (correlation between 99% CI $r_{BWAS}$ and max rankings = 0.96; correlation between 99% and 95% CI $r_{BWAS}$ rankings = 0.99). For cortical thickness data, we correlated each of the 649 variables with each of the 59,412 cortical vertices (fig. S2A).

Multivariate analyses: In addition to univariate associations, we used ridge regression models ($\alpha$ = 1.0), to estimate out-of-sample brain-based (RSFC and cortical thickness, separately) multivariate associations for each of the 649 non-imaging variables (predicted scores vs. observed scores). For input brain features, we first conducted a principal component analysis, whereby the number of principal components retaining 20% of the variance were subsequently passed as features into the ridge regression models. We replicated the ridge regression findings across a range of brain feature sets (number of brain principal components used in training models ranging from 10 - 90% variance explained in the imaging data (*1*)).

*Replication analyses with ComBat and singletons only*

Technical differences between scanners used in the ABCD study could contaminate statistical inference (*108*). Because life factors considered herein may be unevenly distributed across sites and scanners, certain effects could be driven artificially by scanner/site differences. Thus, we harmonized RSFC data using the batch-adjustment algorithm ComBat (*113*), and subsequently correlated each life factor with harmonized data to demonstrate high reproducibility of brain-wide association strength rankings to RSFC data that was not harmonized using ComBat (Supplementary Table 1).

Additionally, the ABCD Study contains siblings and twins. We repeated the correlations of life factor associations with the brain using a sample of only singletons to demonstrate high reproducibility of brain-wide association strength rankings to the sample that also included siblings and twins (Supplementary Table 1).

*Brain-wide association maps*

RSFC network maps of exemplar phenotypes (SES: social and economic domain of the Child Opportunity Index, sleep duration, screen time (total screen time usage during the week and weekend), IQ (NIH Toolbox Cognition Battery, total score) are plotted in Fig. 2A,B. Maps were constructed by averaging the absolute value of association strengths (bivariate r's) across all edges for each region of interest. Each resulting brain network map was subsequently min/max normed, separately within each variable.

Cortical thickness maps for the same 649 variables were generated by projecting the mean bivariate *r* of all cortical vertices within an ROI and variable onto the cortical surface in Workbench Connectome (reference (*114*); fig. S4A.B)). To quantify the similarity between each brain map, we correlated each brain map with every other brain map (at the edge level for RSFC (Fig. 2C) and vertex level for cortical thickness (fig. S4C)), resulting in a 649 x 649 brain map similarity matrix.

*Generation and benchmarking of the principal exposome brain map*

We employed principal component analysis (PCA) across all 649 brain maps (edge x variable for RSFC; vertex x variable for cortical thickness) and extracted the scores from the first principal component, which represents an orthogonal vector containing the greatest amount of variance shared across all brain maps. Brain maps in Fig. 2D (fig. S4D for cortical thickness) were constructed by averaging the absolute value of association strengths (bivariate *r*'s) across all edges for each region of interest (RSFC; all vertices within a region of interest for cortical thickness).

We computed the spatial correlation between each brain-wide association map and the principal brain map and subsequently ranked them by their correlation strength with the exposome brain map (Fig. 2E for RSFC; fig. S4E for cortical thickness).

*Network-level associations*

Regions of interest were ascribed to their respective functional networks (Fig. 2F; fig. S4F for cortical thickness). We performed an independent sample t-test to determine whether the exposome map exhibited significantly stronger associations with sensory and motor networks than all other networks.

*Linear mixed effects modeling of RSFC with socioeconomics, sleep, screen time, and intelligence*

To jointly consider the independent contributions of the strongest life factor BWASs (socioeconomics: Child Opportunity Index, screen time: total screen time, sleep duration, and IQ scores), we used linear mixed-effects models to determine effect sizes of the aforementioned variables. fig. S12A compares resulting standardized beta values from mixed effect models including all variables, as well as those from independent mixed models (for example, socioeconomics modeled separately from sleep duration, screen time, and IQ). All mixed models included fixed effects of age, sex, head motion (mean FD) and random effects of site. fig. S12B-E contains the residualized brain maps for each variable.

*Non-brain variable correlations*

We quantified the correlation (*115*) between each of the non-brain variables using bivariate correlations ($r_{vars}$; fig. S9) and presented exemplar variables strongly related to SES (social and economic domain of the Child Opportunity Index) in fig. S8A.

*Comparative functional pattern analytics*

To derive a functional interpretation to the sensory and motor pattern exhibited by the exposome map, we quantified the similarity of the exposome BWAS map with other strong BWAS maps (SES, IQ), task fMRI maps (working memory, reasoning, and cognitive control) from Neurosynth (*116*), and maps of norepinephrine receptor density from reference (*45*), and stimulants (from trial fMRI (*46*)). We quantified the correlation between these functional brain maps and tested them for significance using spin tests.

To test the hypothesis that the sensory and motor pattern was significantly different between the task fMRI maps (Fig. 3A, left column) and BWAS (Fig. 3A, middle column) and arousal maps (Fig. 3A, right column), we first quantified the proportion of normalized associations that were spatially located in sensory and motor networks (visual, dorsal motor, lateral motor, auditory) vs. all others for each map. Next, we submitted these values for each map into a one-way analysis of variance (ANOVA), grouping by map type (task fMRI, BWAS, arousal). Because the omnibus test was significant, we performed follow-up t-tests to test each pair of map types using the Tukey-Kramer implementation in Matlab.

*Influence of neighborhood and familial socioeconomics on brain with intelligence score associations*

Univariate RSFC with IQ scores (Fig. 4A; NIH Toolbox Cognition Battery, total score) and cortical thickness with IQ scores (fig. S14A) associations were quantified unadjusted for SES, as well as adjusted for SES (social and economic domain of the Child Opportunity Index). All models employed linear mixed effects modeling, covarying for the fixed effects of age, sex, and mean framewise displacement (FD; a composite measure of head motion), as well as a random effect of site. The resulting *t*-statistic was extracted for each univariate brain feature with IQ scores from unadjusted and adjusted models and plotted as a histogram (Fig. 4A).

We observed that 95% of univariate RSFC with IQ score associations were either no longer significant (68%; $P > 0.001$, FDR corrected) or inflated when adjusting for SES. To determine whether the brain features (RSFC: edges; cortical thickness: vertices) decreased in a systematic or random manner in their association with IQ scores, we correlated the change in *t*-statistics in SES adjusted vs. unadjusted models (Fig. 4B, y-axis; fig. S14B, y-axis for cortical thickness) with the *t*-statistics for brain with IQ in unadjusted models (Fig. 4B, x-axis; fig. S14B, x-axis for cortical thickness).

*Adjusting multivariate associations of brain with IQ for SES*

We used ridge regression to assess out-of-sample multivariate association between the brain and IQ scores (NIH Toolbox Cognition Battery, total score). Data were randomly split into two equal halves - a training set and test set - across the full sample (Discovery and Replication combined; $n$ = 4,579) and the following steps were repeated across 100 bootstrapped samples. First, principal component analysis was conducted on the training set RSFC data. Components explaining 20% of between-person variance in RSFC (34 components) were extracted and submitted to the training model. The test set RSFC data were projected into the principal component coordinate space of the training RSFC data to avoid leakage. The intercept and beta weights were applied to the testing RSFC data to produce "predicted" IQ scores. These scores were correlated with the observed IQ scores in the test set, resulting in the out-of-sample multivariate association presented in Fig. 4C for RSFC and Supplemental Fig. 8C for cortical thickness. We chose 20% of brain features to balance data reduction with maximizing out-of-sample (replication) association strength, a hyperparameter choice made in previous work and carried forward here to mitigate model overfitting . Moreover, our prior work(*1*) has shown that reproducible multivariate brain-behavior associations using similar CCA models are maximized by including ~20% of RSFC components.

We repeated the above procedure after regressing SES (social and economic domain of the Child Opportunity Index) from brain components in the Replication (test) dataset. In Fig. 4C, we report and compare the out-of-sample multivariate association for RSFC (fig. S14C for cortical thickness) with IQscore models adjusting for SES with those unadjusted for SES.

*Adjusting multivariate associations of brain with SES for IQscores*

We used ridge regression to assess out-of-sample multivariate association between the brain and SES (social and economic domain of the Child Opportunity Index). Data were randomly split into two equal halves - a training set and test set - across the full sample (Discovery and Replication combined; $n$ = 4,579) and the following steps were repeated across 100 bootstrapped samples. First, principal component analysis was conducted on the training set RSFC data. Components explaining 20% of between-person variance in RSFC (34 components) were extracted and submitted to the training model. The test set RSFC data were projected into the principal component coordinate space of the training RSFC data to avoid leakage. The intercept and beta weights were applied to the testing RSFC data to produce "predicted" SES scores. These scores were correlated with observed SES in the test set, resulting in the out-of-sample multivariate association presented in fig. S17A for RSFC and fig. S17B for cortical thickness.

We repeated the above procedure after regressing IQ (NIH Toolbox Cognition Battery, total score) from brain components in the Replication (test) dataset. In fig. S17A, we report and compare the out-of-sample multivariate association for RSFC (fig. S17B for cortical thickness) with SES models adjusting for IQ scores with those unadjusted for IQ scores.

*Connectome-based prediction model: Multivariate associations of brain with IQ*

We implemented the connectome-based prediction model (*55*) (CPM) to generate an out-of-sample association between brain function (RSFC; fig. S15) and IQ scores (NIH Toolbox Cognition Battery, total score; fig. S16 for cortical thickness), adjusted for SES (social and economic domain of the Child Opportunity Index) and unadjusted. Out-of-sample association strength ($r_{oos}$) is reported as the correlation between predicted and observed phenotypic scores (using models trained on the Discovery set). We repeated this analysis 100 times on random split-halves of the data. SES was regressed from the observed test set IQ scores prior to correlating predicted vs. observed values for adjusted models (*57*).

*Additional deconfounding approaches in multivariate modeling*

Regressing confounds from both the training and test datasets can result in either inflated or deflated out-of-sample associations and is not recommended (*57*). For the multivariate models presented in Fig. 4 and fig. S17, in which we controlled for SES when predicting IQ scores from either RSFC or cortical thickness data, we regressed SES in the test sample only. Similarly, in brain-based models of SES, we regressed IQ scores from the test set.

We also performed analyses of multivariate brain-based associations with IQ scores and SES, separately, using a broader form of a confound isolation cross-validation procedure (*57*). We conducted this set of analyses using ridge regression, given a single target phenotype (either IQ scores or SES).

In confound isolation cross-validation (*57*), a test set of data is generated from the full dataset such that the confound, "z" (in this caseSES; social and economic domain of the Child Opportunity Index) and the target, "y", (in this case IQ; NIH Toolbox Cognition Battery, total score) are uncorrelated. Given the strong correlation between SES and IQ scores in the ABCD data ($r_{vars}$ = 0.31), it is difficult to generate random samples in which the correlation between SES and IQ are close to 0 at larger samples. To generate such samples, we relied on a nested cross validation framework consisting of a test set of *n* = 50. Across 1,000 iterations, we randomly selected test samples of *n* = 50 and extracted the correlation between SES and IQ (fig. S18). Brain-based (RSFC) models of IQ were trained on the remaining *n* = 4,529 participants and then tested on the *n* = 50 test sample.

*Generalizability of multivariate brain with IQ associations*

For all generalizability analyses, we used canonical correlation analysis (CCA), a common multivariate technique used to learn associations between multivariate brain and non-brain behavioral data, carrying forward hyperparameters from our previous work(*1*) to reduce overfitting. CCA learns associations between multivariate brain and non-brain data. Both IQ scores (NIH Toolbox Cognition Battery, total scores) and SES (social and economic domain of the Child Opportunity Index) are composite measures composed of multiple subscales. For all generalizability analyses in Fig. 5,6, fig. S19, fig. S22 to S24, fig. S26 to S28. Generalizability was assessed by training models using the ABCD Discovery dataset and testing models on the ABCD Replication sample and the HCP dataset.

We tested the out-of-sample generalizability of multivariate models of RSFC and cortical thickness (separately) with IQ scores (NIH Toolbox Cognition Battery, all subscales) in subsamples of varying SES composition. For these analyses (RSFC: Fig. 5; cortical thickness: fig. S19), we trained models using the ABCD data containing subsamples (*n* = 569) of individuals from only either the full Discovery sample, individuals from higher SES (z-scored social and economic domain of the Child Opportunity Index > 0.75) or lower SES (z-scored social and economic domain of the Child Opportunity Index < -0.25). In each case, the IQ distributions of the training (Discovery) and test (Replication) set were matched.

We also tested for generalizability of brain-based IQ models using CPM. For CPM analyses, we trained models using the ABCD data containing individuals from only either lower socioeconomic opportunity (z-scored social and economic domain of the Child Opportunity Index < 0.55, which includes *n* = 569 individuals or higher socioeconomic opportunity (z-scored social and economic domain of the Child Opportunity Index > 0.75, which includes *n* = 529 individuals > 0.5 standard deviations from the median z-scored social and economic domain of the Child Opportunity Index value of 0.25). Because CPM cannot learn associations between two multivariate dataset, we used IQ scores (NIH Toolbox Cognition Battery, total score) as the target variable. Each model was subsequently tested separately on (i) all individuals in the replication dataset (full), (ii) individuals from lower socioeconomic opportunity neighborhoods (low), and (iii) individuals from high socioeconomic opportunity neighborhoods (high). This 2 x 3 design was run for CCA models of RSFC with IQ scores (fig. S20), cortical thickness with IQ scores (fig. S21).

*Shortcut learning in brain-based models of IQ*

We tested the out-of-sample generalizability of multivariate models (CCA) of RSFC with IQ scores (NIH Toolbox Cognition Battery, all subscales) for varying levels of SES using the ABCD Discovery dataset (Fig. 6A) as a training dataset and either the ABCD Replication dataset (Fig. 6a) or the HCP (adult) dataset (fig. S26) as a test dataset. For supplemental analyses on cortical thickness, see fig. S22. We trained models using the ABCD data containing varying subsample compositions with regards to SES (*z*-scored social and economic domain of the Child Opportunity Index), ranging from high (containing only high SES individuals) to full sample (containing individuals randomly sampled from all levels of SES), with a total of 54 bins that were bootstrap resampled 100 times for a total of 5,400 models. Each ABCD training set consisted of $n = 569$ individuals for testing on the ABCD Replication dataset. For training models tested on the HCP sample, $n = 877$ individuals were used to train models, in order to match the HCP sample size. The mean across the 100 resamples +/- one standard deviation is plotted in Fig. 6A and fig. S26. For each of the 5,400 models, we quantified the correlation between IQ, the target variable, and SES (Fig. 6B).

*Negative control*: We repeated the above analyses regarding the generalizability of multivariate models (CCA) of RSFC with IQ scores within the ABCD dataset, but rather than varying subsample composition with respect to SES in the ABCD training sample, we varied the level of total psychopathology scores (CBCL total score; p-factor) from low (indicates no psychopathology), to full sample (individuals randomly sampled from the whole p-factor distribution). Each ABCD training (Discovery) and test (Replication) set consisted of $n = 569$ individuals. The mean across the 100 resamples +/- one standard deviation also is plotted in Fig 6a.

*No Shortcut learning in brain-based models of SES*

We tested the out-of-sample generalizability of multivariate models (CCA) of RSFC with SES (social and economic domain of the Child Opportunity Index, all subscales) for varying levels of IQ ($80 < IQ < 110$) using the ABCD Discovery dataset as a training dataset and the ABCD Replication dataset (Fig. 6c) as a test dataset. We trained models using the ABCD data containing varying subsample compositions levels of IQ (NIH Toolbox Cognition Battery, total score), ranging from high IQ only (containing only individuals with IQ > 110) to the full sample (containing individuals randomly sampled from all levels of IQ), with a total of 31 bins that were bootstrap resampled 100 times for a total of 3,100 models. Each ABCD training set consisted of $n = 569$ individuals for testing on the ABCD Replication dataset. The mean across the 100 resamples +/- one standard deviation is plotted in Fig. 6C and fig. S24A.

*Negative control*: We repeated the above analyses regarding the generalizability of multivariate models (CCA) of RSFC with SES within the ABCD dataset, but rather than varying subsample composition with respect to IQ in the ABCD training sample, we varied the level of total psychopathology scores (CBCL total score; p-factor) from low (indicates no psychopathology), to full sample (individuals randomly sampled from the whole p-factor distribution). Each ABCD training (Discovery) and test (Replication) set consisted of *n* = 569 individuals. The mean across the 100 resamples +/- one standard deviation also is plotted in Fig. 6C and fig. S24A.

<u>*Genetic ancestry*</u>: We tested the out-of-sample generalizability of multivariate models (CCA) of RSFC with SES (social and economic domain of the Child Opportunity Index, all subscales) within the ABCD dataset, varying the training sample by genetic ancestry (genetic principal components, of which there were 32 components; the most restrictive sample composition Fig. 9a, left) indicates greater homogeneity with respect to genetic ancestry along a given component, whereas “all” indicates individuals randomly sampled from the whole ancestral distribution). We varied the training sample by each genetic principal component separately. For fig. S24A, we first averaged the out-of-sample correlation ($r_{oos}$) across the 100 bootstrapped samples within a genetic principal component and subsequently averaged over all 32 genetic principal components. Each ABCD training (Discovery) and test (Replication) set consisted of *n* = 569 individuals. The mean across the 32 genetic principal components +/- one standard deviation is plotted in fig. S24A. For out-of-sample correlations ($r_{oos}$) when varying the training sample for each genetic principal and for each bootstrapped sample, see fig. S25.

<u>*Generalizability of brain with IQ score and SES associations across sexes*</u>

To demonstrate specificity of lower generalizability of brain connectivity with IQ score associations as a function of SES, we trained RSFC with IQ models and RSFC with SES models (separately) using either females or males only in the Discovery dataset. Subsequently, we tested each of those models in the replication dataset of either female only or male only (fig. S27). We repeated this analysis for brain connectivity with socioeconomic (social and economic domain of the Child Opportunity Index, all subscale) associations (fig. S28).

<u>*Significance testing*</u>

Significance testing included both parametric and nonparametric tests. We keep with scientific standards in this regard, including reporting of corrected *P* values where appropriate. We report all Discovery and Replication effect sizes in the tables and figures to encourage interpretation of effect sizes over *P* values when using large (*n* > 1,000) population-level datasets.

#### Supplementary Tables

**Supplementary Table 1. List of ABCD variables and effect sizes**
Please see the attachment.

**Supplementary Table 2. Receptor map correlations with SES brain map**

| Receptor map | Correlation with SES map ($r_{patterns}$) |
|---|---|
| Norepinephrine | 0.35 |
| Glutamate (NMDA) | 0.15 |
| GABAa | 0.08 |
| Acetylcholine | 0.07 |
| Glutamate (R5) | 0.03 |
| Dopamine (D2) | 0.00 |
| Serotonin (5HT2a) | -0.11 |
| Cannabinoid | -0.12 |
| Dopamine (D1) | -0.17 |
| Serotonin (5HT1a) | -0.31 |
| Opioid | -0.35 |

#### Supplementary Text

##### Socioeconomic permeation of brain-wide associations

The brain-wide association map similarity to SES was stronger for some variables (sleep duration, screen time, IQ scores ($r_{patterns} > 0.70$; Fig. 2B) than others (psychopathology (p-factor); $r_{patterns} = 0.29$; Fig. 2B). A straightforward explanation could be that some of the differences in association map similarities arise from the correlations between the underlying non-imaging variables ($r_{vars}$).

Indeed, variables with univariate brain-wide association patterns similar to SES also had strong direct variable-to-variable correlations with SES (fig. S8; IQ scores $r_{vars} = 0.31$, $P = 6.78 \times 10^{-53}$; sleep duration $r_{vars} = 0.30$, $P = 1.27 \times 10^{-50}$; screen time $r_{vars} = -0.24$, $P = 2.70 \times 10^{-31}$), while the direct correlation between psychopathology (p-factor) and SES was weaker (fig. S8; $r_{vars} = -0.09$, $P = 3.28 \times 10^{-5}$; fig. S9 for full $r_{vars}$ correlation matrix). To evaluate the relative association strength of SES, sleep duration, screen time, and IQ scores with brain connectivity, we combined them in the same multiple regression model. This did not alter their relative ranking, as SES remained the strongest association (fig. S12A; $P < 0.0001$).

The SES sensory/motor cortex association pattern permeated many other BWAS maps (Fig. 2), raising the question of whether SES similarly contributes to BWAS association strengths of non-socioeconomic variables (99% CI $r_{BWAS}$). Therefore, we plotted BWAS association strengths ($r_{BWAS}$; fig. S8, y-axis), for all non-socioeconomic variables, against their variable-to-variable correlation ($r_{vars}$) with childhood SES (fig. S8, x-axis). Across all non-socioeconomic variables, BWAS association strengths (fig. S8; y-axis: $r_{BWAS}$ 99% CI for each variable) and correlations with SES (fig. S8; x-axis: $r_{vars}$) were strongly related ($r = 0.87$, $P = 6.72 \times 10^{-198}$; fig. S8; fig. S10 for cortical thickness). This relationship suggests that some BWAS (for example, IQ scores) are stronger than others (for example, p-factor) because of their stronger variable-to-variable correlation with SES (fig. S8; IQ scores $r_{vars} = 0.31$; psychopathology $r_{vars} = -0.09$). However, some variables (sleep duration, screen time) had even stronger brain associations ($r_{BWAS}$; fig. S8; y-axis) than would be expected by their correlation ($r_{vars}$; x-axis) with SES (fig. S8; above the fit line), while other variables showed weaker than expected associations based on their relationship with SES (neighborhood safety/crime; fig. S8; below the fit line). Variables substantially above the best fit line (fig. S8, black), such as sleep duration and screen time, are associated with child brain function beyond their relationship with SES. Interestingly, variables associated with arousal-related brain function (e.g., sleep duration) were not substantially above the best-fit line for cortical thickness (fig. S10).

### Supplementary Figures

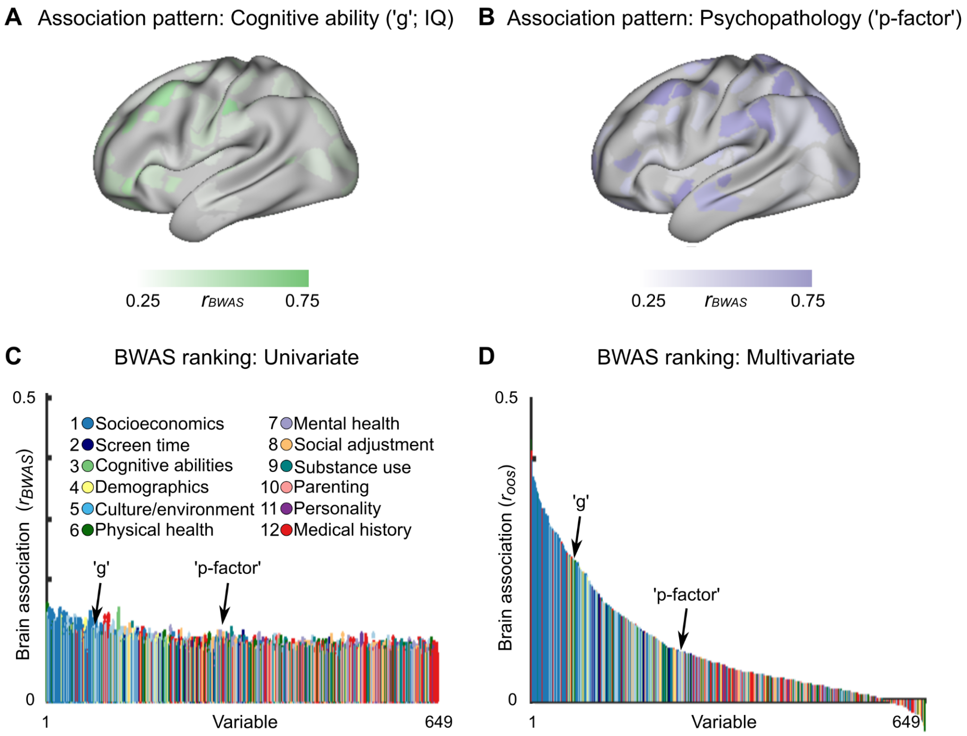


**fig. S1. Rank order of cortical thickness brain-wide associations across 649 variables.** Cortical thickness BWAS maps (bivariate correlation, $|r_{BWAS}|$) for commonly studied exemplar variables: **(A)** cognitive ability ('g', IQ scores; NIH Toolbox Cognition Battery, total score) and **(B)** psychopathology ('p-factor', Child Behavior Checklist, total score) on a predefined Discovery ($n$ = 2,350; training) set of Baseline ABCD data. **(C)** Brain-behavior association strength (y-axis) for 649 non-imaging variables (x-axis). Each colored dot (59,412 dots for each variable) represents a single association between a cortical thickness vertex and a non-imaging variable. Dot color represents the predefined category of the variable. The non-imaging variables (x-axis) are ranked by their 99% confidence interval across all 59,412 associations. All associations were based on at least $n$ > 2,000 individuals. **(D)** Multivariate brain-behavior association (bivariate correlation ($r_{oos}$) between out-of-sample predicted and observed scores) of brain-based cortical thickness models for each non-brain variable using ridge regression (see Methods). Multivariate brain models were trained using ridge regression on a predefined Discovery ($n$ = 2,350; training) set of Baseline ABCD data and subsequently tested on a matched (see Methods) left-out Replication ($n$ = 2,276; test) set of Baseline ABCD data. Dot color represents the predefined category of the variable. The non-imaging variables (x-axis) are ranked by their multivariate association strength ($r_{oos}$).

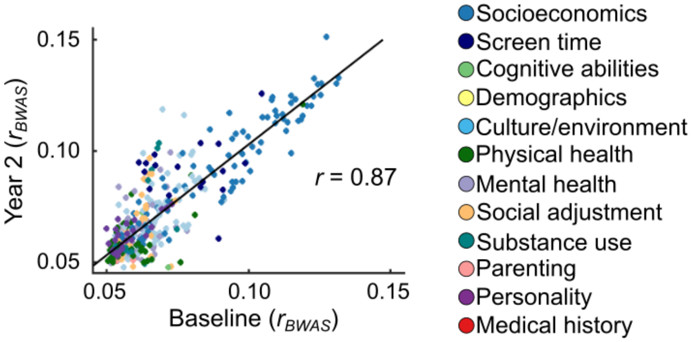


**fig. S2. Correlation of univariate effect sizes between ABCD Baseline and Year 2 waves.** Year 2 (y-axis; *n* = 2,363) vs. Baseline (x-axis; Discovery sample: *n* = 2,316; 99$^{th}$ percentile (confidence interval)) univariate resting-state functional connectivity (RSFC) effect sizes (99$^{th}$ percentile (confidence interval) of univariate effect sizes ($r_{BWAS}$) for each non-brain variable) in the ABCD Study. The high correlation between Baseline and Year 2 ($r$ = 0.87) indicates stable effect size rankings across study waves. Dots represent a given RSFC variable pair ($r_{BWAS}$), and the colour of the dot denotes the category to which the variable belongs. Black line represents the line of best fit.

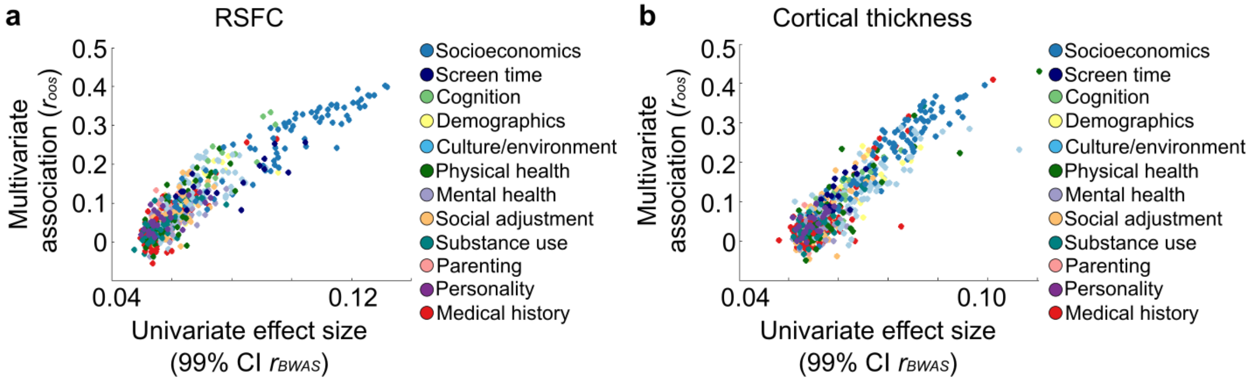


**fig. S3. Relationship of multivariate and univariate associations. (a)** Correlation between multivariate association strengths prediction accuracy (y-axis; out-of-sample bivariate correlation (*$r_{oos}$)* between predicted scores and observed variable scores) using ridge regression (see Methods) and univariate association strengths (99$^{th}$ percentile (confidence interval)top 1% bivariate associations between resting-state functional connectivity (RSFC) and each non-brain variable) in the Baseline ABCD sample across all variables. Multivariate models were trained on the Baseline ABCD Discovery sample (*n* = 2,316) and subsequently tested on the Replication sample (*n* = 2,263). The color of the dot represents the predefined category of the variable. **(b)** Correlation between multivariate association strengths prediction accuracy (y-axis; out-of-sample bivariate correlation (*$r_{oos}$)* between predicted scores and observed variable scores) using ridge regression (see Methods) and univariate association strengths (99$^{th}$ percentile (confidence interval)top 1% bivariate associations between cortical thickness and each non-brain variable) in the Baseline ABCD sample across all variables. Multivariate models were trained on the Baseline ABCD Discovery sample (*n* = 2,351) and subsequently tested on the Replication sample (*n* = 2,276). The color of the dot represents the predefined category of the variable.

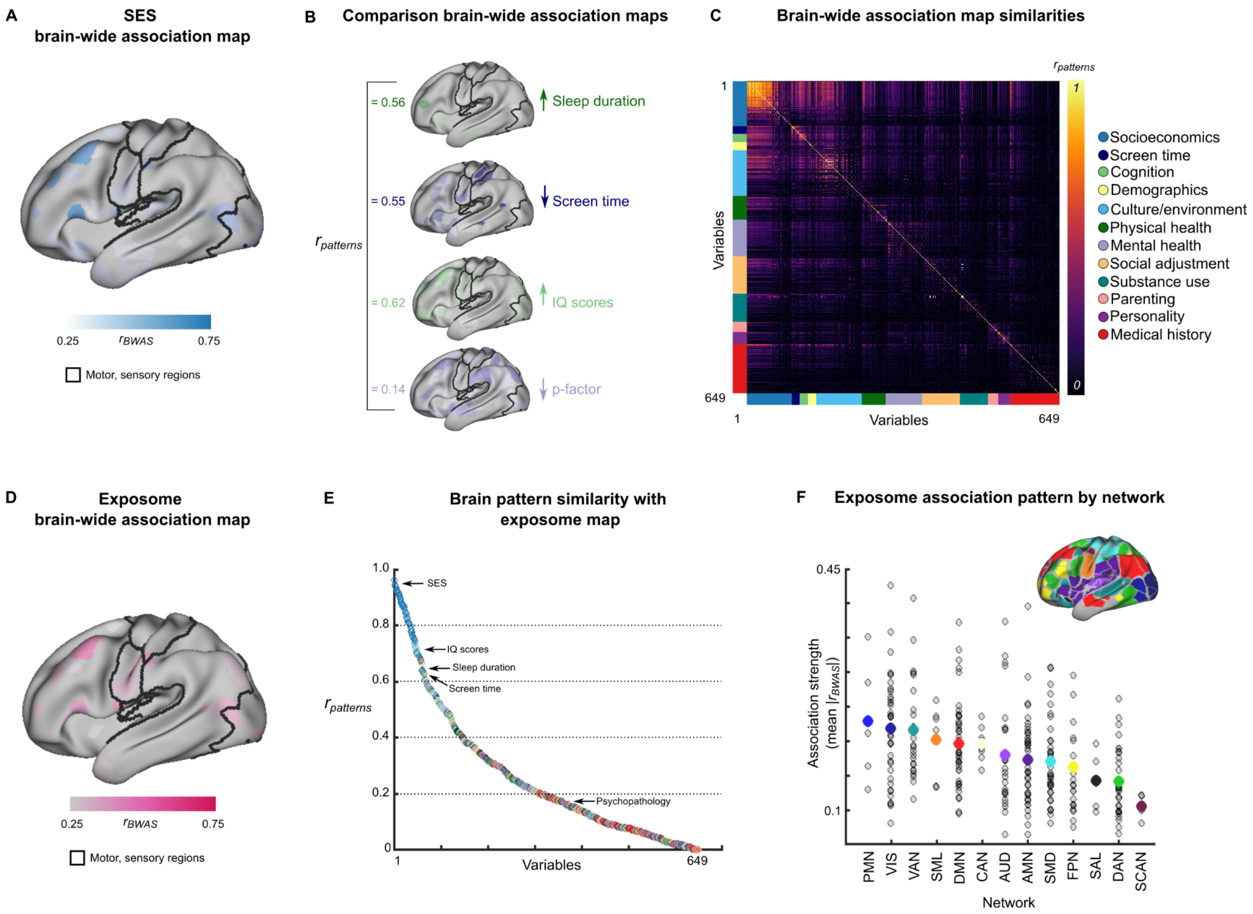


**fig. S4. Brain-wide association map comparisons for cortical thickness. (A)** Brain-wide association map of cortical thickness with SES (social and economic domain of the Child Opportunity Index; see fig. S6A for medial wall and right hemisphere). Values are min/max (0-1) normed. **(B)** Exemplar variables for comparison (fig. S6 for additional brain views). The bivariate correlations ($r_{patterns}$) between the brain-wide association map for SES (panel **A**) and comparison brain-wide association maps (panel **B**) are listed in the left column of panel **B**. Spin tests revealed significant associations between BWAS maps for SES, sleep duration, screen time, and IQ (NIH Toolbox Cognition Battery, total score, $P < 0.001$, Bonferroni corrected). **(C)** Similarity matrix depicting the correlations ($r_{patterns}$) for all possible pairs of brain-wide association maps (649 x 649). Variables are organized according to category (right-sided legend) and sorted by association strength within each category. Color tabs on the x- and y-axes reflect the category to which a variable belongs. **(D)** The exposome association map is the first principal component across all 649 brain-wide association maps. See fig. S6 for right hemisphere, medial views. **(E)** Correlations ($r_{patterns}$; y-axis) of the brain-wide association maps for each variable (x-axis) with the exposome association map (from panel **E)**. Variables are ranked according to their spatial correlation with the exposome brain-wide association map. The right side of the line break on the x-axis shows the mean +/- the standard error for each category. **(F)** Association strength (mean normed $|r_{BWAS}|$; y-axis) of the exposome map from **D** for each functional brain network (x-axis; see Methods). Each grey circle represents the mean association strength ($|r_{BWAS}|$) for a parcel/region. The coloured dot for each network represents the mean association strength across all regions within that network. SMD: sensorimotor dorsal; VIS: visual; SCAN: somato-cognitive action; SML: somatomotor lateral; AUD: auditory; VAN: ventral attention; AMN: action mode; DMN: default mode; DAN: dorsal attention; CAN: context association; PMN: parietal memory; SAL: salience; FPN: frontoparietal.

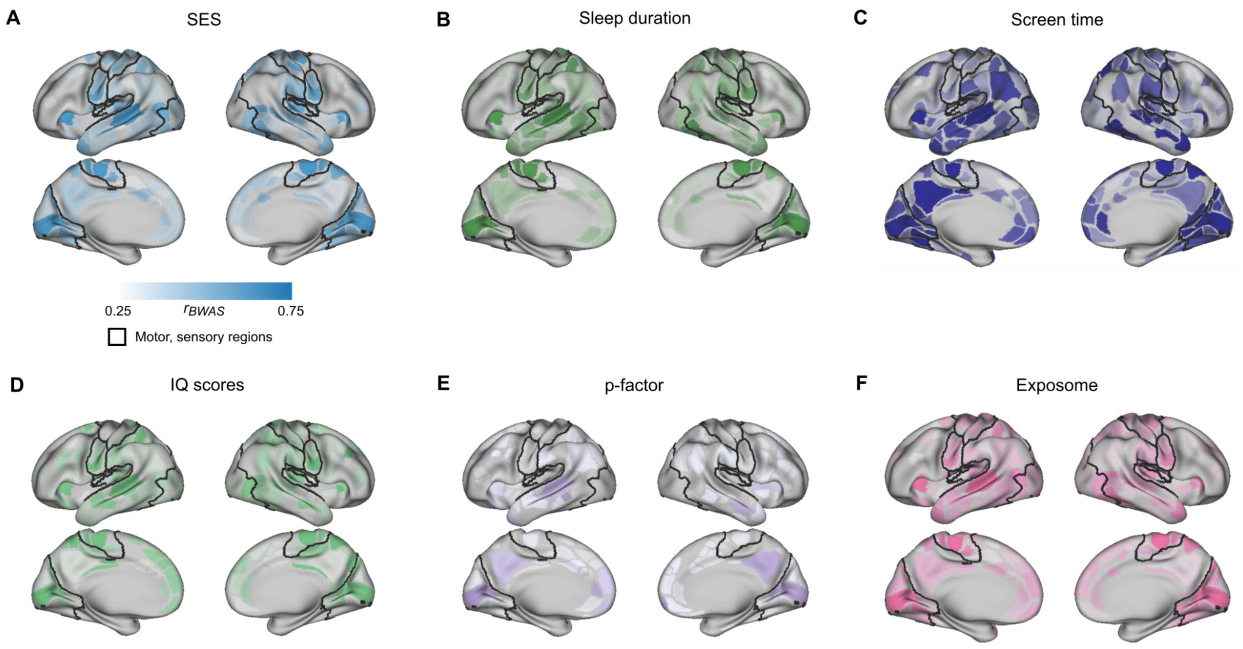


**fig. S5. Resting-state functional connectivity brain-wide association maps additional views.** Associations between resting-state functional connectivity (RSFC) and variables from Fig. 2 (ABCD), including **(a)** SES (the social and economic domain of the Child Opportunity Index) **(b)** sleep duration, **(c)** total screen time, **(d)** IQ scores (NIH Toolbox Cognition Battery, total score), **(e)** p-factor (total score on the Child Behavior Checklist (CBCL)) and **(f)** the exposome brain map (that is, the first principal component across all 649 variable brain maps). For display purposes, each parcel is colored according to the sum of the absolute value of association strengths ($r_{BWAS}$) from that parcel to every other parcel. Each brain map is min/max normed, with deeper hues representing parcels with relatively stronger associations ($r_{BWAS}$) with the variable of interest.

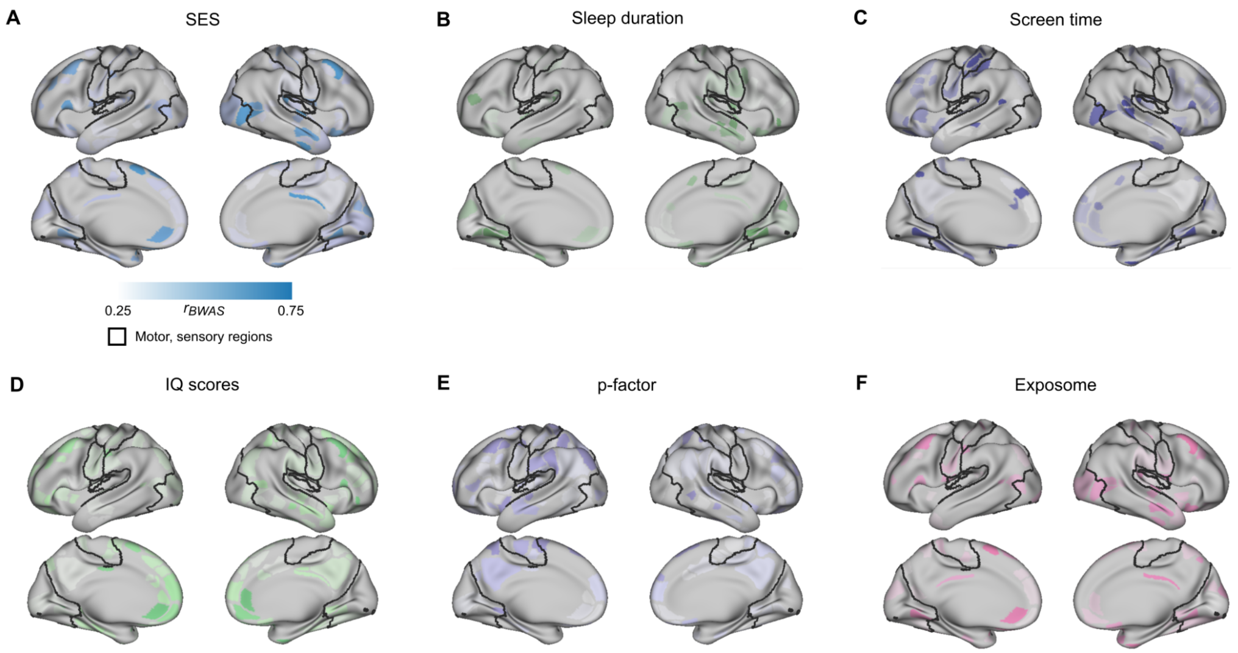


**fig. S6. Cortical thickness brain-wide association maps.** Associations between cortical thickness and variables from fig. S4 (ABCD), including **(a)** SES (the social and economic domain of the Child Opportunity Index) **(b)** sleep duration, **(c)** total screen time, **(d)** IQ scores (NIH Toolbox Cognition Battery, total score), **(e)** p-factor (total score on the Child Behavior Checklist (CBCL)) and **(f)** the exposome brain map (that is, the first principal component across all 649 variable brain maps). For display purposes, each parcel is colored according to the normalized (min/max) sum of the absolute value of association strengths ($r_{BWAS}$) from that parcel to every other parcel. Each brain map is min/max normed, with deeper hues representing parcels with relatively stronger associations ($r_{BWAS}$) with the variable of interest.

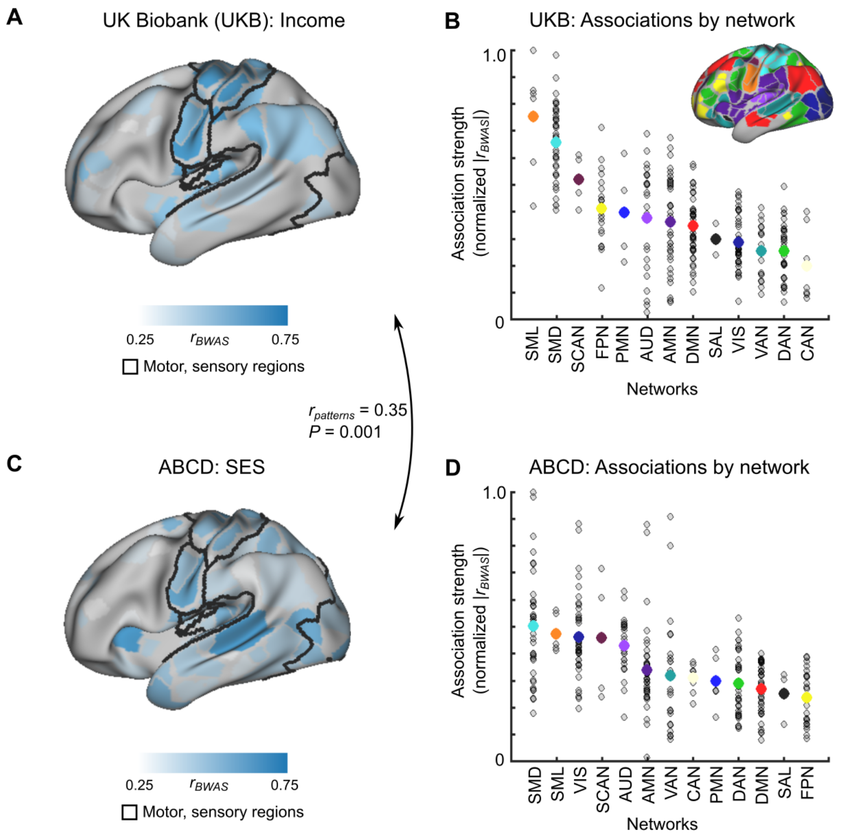


**fig. S7. Univariate associations between resting-state functional connectivity (RSFC) and SES in the UK Biobank and ABCD data sets**. **(A)** Brain-wide association map between RSFC and income in the UK Biobank ($n$ = 32,572, 95% white British, white Irish, or other White background). Blue colors represent brain regions with relatively stronger associations between RSFC and income. **(B)** Association strength by functional network. Similar to ABCD, the somatomotor networks demonstrated the strongest RSFC with income associations. Network template is in the top right quadrant. Values are min/max normalized across 333 brain regions. **(C)** Brain-wide association map between RSFC and SES (social and economic domain of the Child Opportunity Index) in the ABCD Discovery sample ($n$ = 2,316). Blue colors represent brain regions with relatively stronger associations between RSFC and socioeconomic opportunity. **(D)** Association strength by functional network. Values are min/max normalized across 333 brain regions.

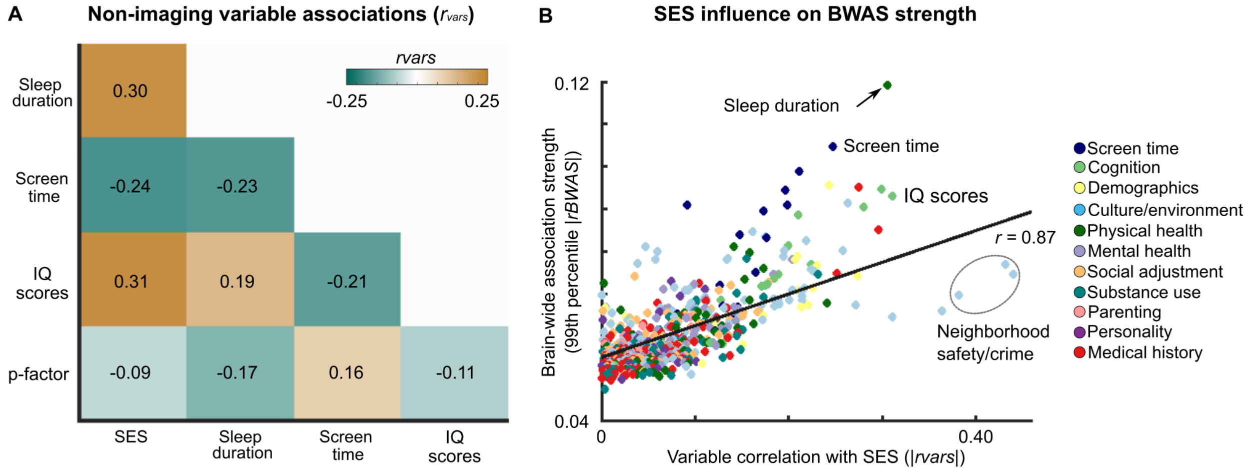


**fig. S8. Associations between SES and other variables. (A)** Correlations between the key non-imaging variables ($r_{vars}$) shown in Fig. 2C, as well as the p-factor. See fig. S9 for the full correlation matrix of 649 non-imaging variables. **(B)** Univariate associations (99% confidence interval from Fig. 1A) of non-socioeconomic variables with brain connectivity ($|r_{BWAS}|$; y-axis) as a function of their correlation with SES ($r_{vars}$; social and economic domain of Child Opportunity Index; x-axis). Variables above the linear best fit line (black line: $r = 0.87$; for example sleep duration, screen time) exhibited stronger associations with brain connectivity than would be predicted by their correlation with SES. See fig. S10 for cortical thickness. Each dot in panel **(B)** depicts a single variable, colored according to its category.

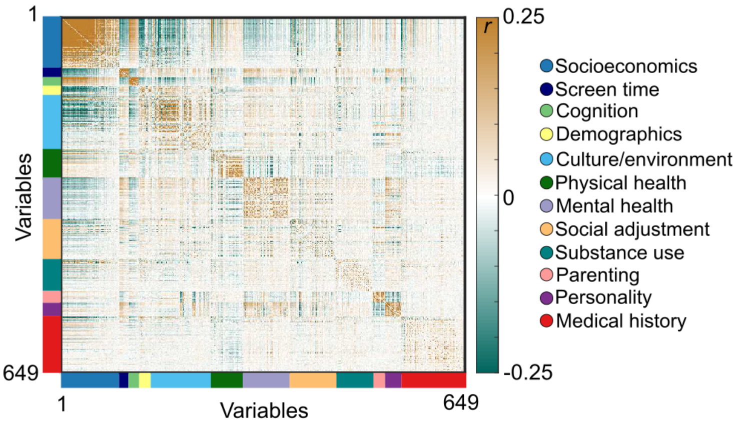


**fig. S9. Correlations between non-brain variables (*$r_{vars}$*).** Correlations (bivariate *r*) between each of the 649 non-imaging variables in the Baseline Discovery ABCD Study dataset (*n* = 2,316). Variables were first sorted by category and subsequently sorted by association strength with resting-state functional connectivity within each category (identical sorting to Fig. 2C). Color bars on the left and bottom axes represent the variable category.

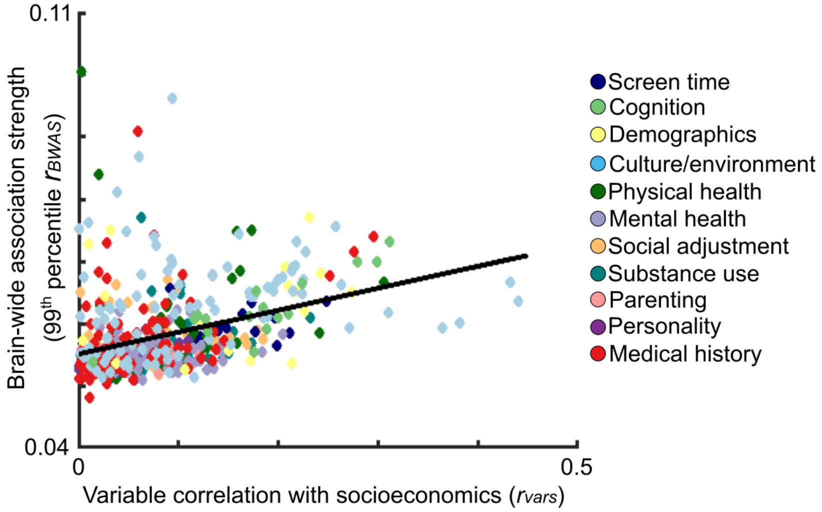


**fig. S10. Associations between SES and other variables for cortical thickness.** Associations (99th percentile (confidence interval) of non-SES variables with cortical thickness ($r_{BWAS}$; y-axis) as a function of their correlation with socioeconomics ($r_{vars}$; social and economic domain of Child Opportunity Index; x-axis). Each dot depicts a single variable, colored according to its category. Variables above the linear best fit line (black line: $r$ = 0.70) exhibited stronger associations with cortical thickness than would be predicted by their correlation with socioeconomics.

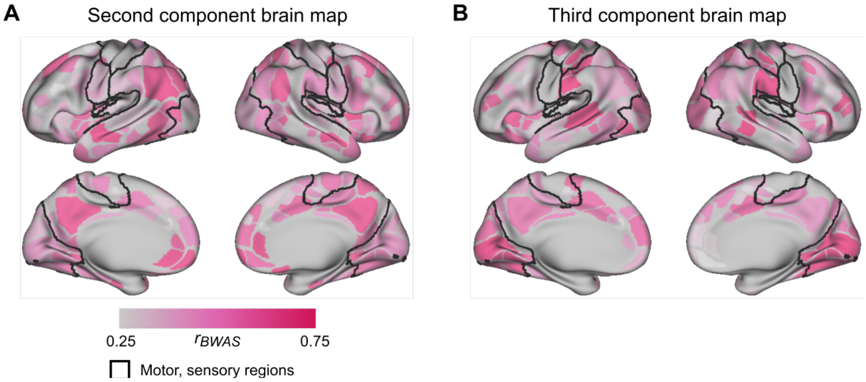


**fig. S11. Brain-wide association maps for additional principal component brain maps. (A)** Second and **(B)** third principal component brain maps (for first principal component brain map, see Fig. 2D). Principal component exposome analysis was run on all 649 brain maps of resting-state functional connectivity (RSFC) with each non-imaging variable. Each brain map is min/max normed, with orange and yellow hues representing parcels with relatively stronger weightings.

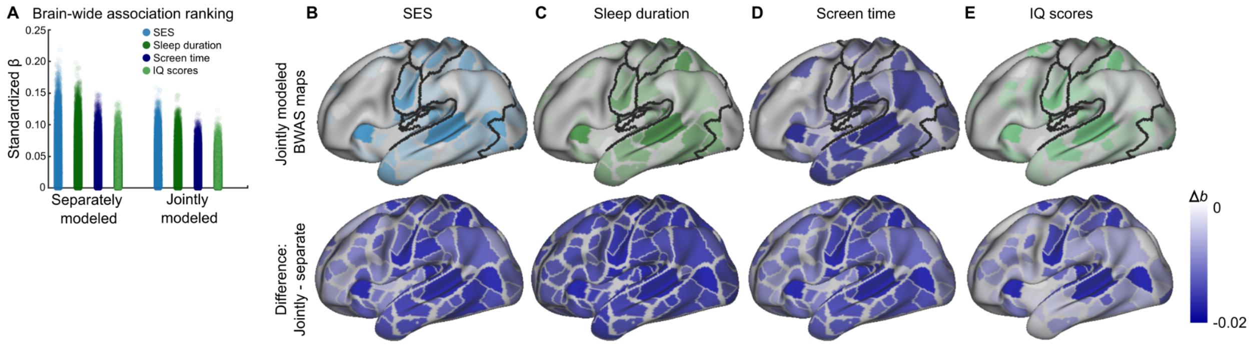


**fig. S12. Residualized brain-wide association maps for functional connectivity. (A)** Brain-wide association strengths (standardized *b;* 99% confidence interval (CI)) for the strongest variables (socioeconomics, sleep duration, screen time, and intelligence scores) in which each variable was considered independently (*left*) and jointly (*right, top*). Brain-wide association strengths are reported here as standardized beta values, since every model was executed using linear mixed effects models (see Methods). Brain-wide association maps of residualized associations (from joint modeling) between resting-state functional connectivity and **(B)** SES (social and economic domain of the Child Opportunity Index), **(C)** sleep duration, **(D)** screen time, and **(E)** IQ scores. The bottom row of panel **b** shows the difference between beta values for each parcel when variables were modeled jointly vs. separate (that is, jointly - separate), showing the brain-wide decrease in association strength.

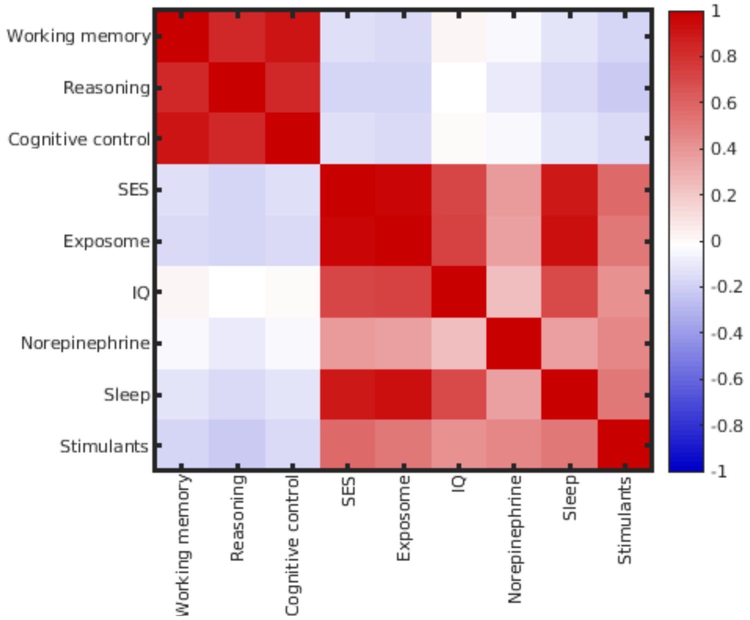


**fig. S13. Correlations between task, BWAS, and arousal brain maps.** Correlations ($r_{patterns}$) between task (working memory, reasoning, and cognitive control) maps from Neurosynth, brain-wide association (BWAS) maps, and arousal-related (norepinephrine, sleep, stimulants) brain function.

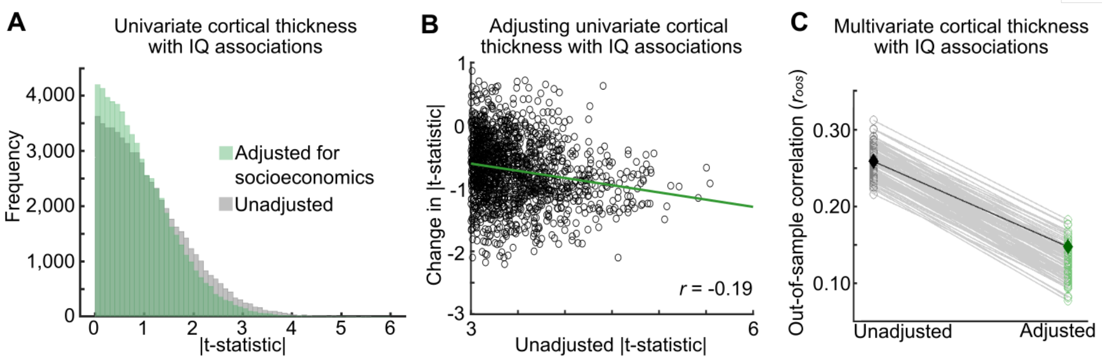


**fig. S14. Adjusting for SES when associating IQ scores with cortical thickness. (A)** Distribution of brain (cortical thickness vertices) with IQ scores (NIH Toolbox Cognition Battery, total score) associations in univariate models unadjusted for SES (grey) and after adjusting for SES (green; adjusting for social and economic domain from the Child Opportunity Index). **(B)** Univariate associations of brain structure (cortical thickness) with IQ scores after SES adjustment; (y-axis) as a function of the unadjusted association between RSFC and IQ scores (x-axis). The negative slope ($r$ = -0.19) indicates that adjusting for SES reduces stronger cortical thickness with IQ associations the most. **(C)** Multivariate associations (out-of-sample correlations: $r_{oos}$) of cortical thickness with IQ scores using ridge regression (See fig. S16 for replication with Connectome-Based Predictive Modeling (*55*)) unadjusted (grey; *left*) for SES and adjusted for SES (green; *right*). Models were trained and tested on 100 split-half bootstrap samples. Black diamond (left) represents the out-of-sample model fit from a predefined replication sample (see Methods), unadjusted for SES, while the dark green diamond (right) represents the out-of-sample correlation from the same predefined matched replication sample (see Methods), adjusted for SES. In all instances, the correlation between observed and predicted scores is plotted.

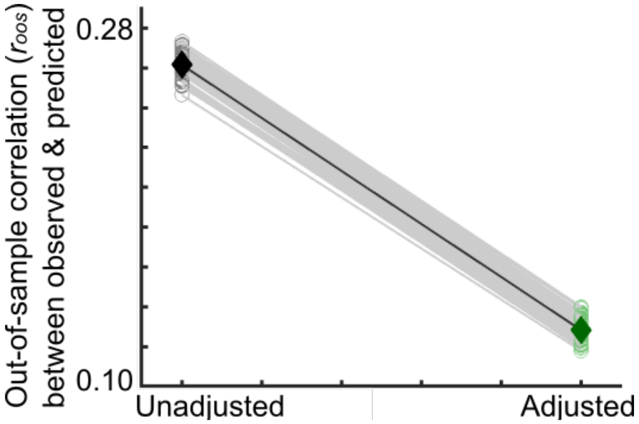


**fig. S15. SES influence on Connectome-based Prediction Model (CPM) associations between IQ scores and functional connectivity.** Multivariate associations of intelligence scores from brain connectivity using the connectome-based prediction model (CPM) unadjusted (grey; *left*) for socioeconomics (normed social and economic composite from the Child Opportunity Index) and adjusting (green; *right*) for socioeconomics. Models were trained and tested on 100 split-half bootstrapped samples. Black dot (left) represents the in-sample model fit from a predefined discovery sample (see Methods), and the dark green dot (right) represents the out-of-sample correlation from a predefined replication sample (see Methods).

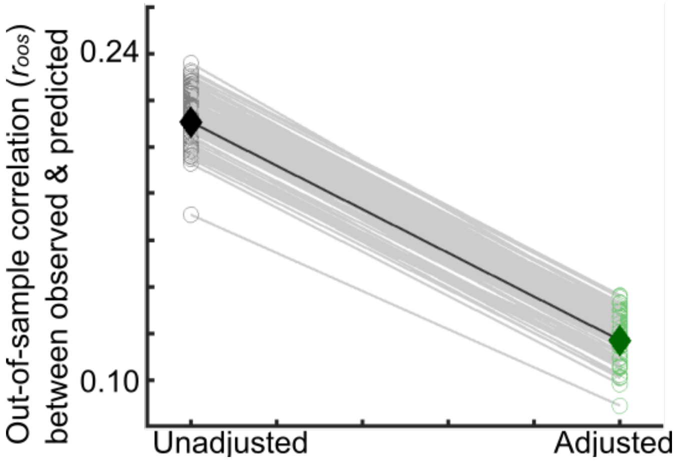


**fig. S16. Socioeconomic influence on Connectome-based Predictive Modeling (CPM) associations between IQ scores and cortical thickness. a)** Multivariate associations of intelligence scores from cortical thickness using the connectome-based prediction model (CPM) unadjusted (grey; *left*) for socioeconomics (social and economic composite from the Child Opportunity Index) and adjusting (green; *right*) for socioeconomics. Models were trained and tested on 100 split-half bootstrapped samples. Black dot (left) represents the in-sample model fit from a predefined discovery sample (see Methods), and the dark green dot (right) represents the out-of-sample correlation from a predefined replication sample (see Methods).

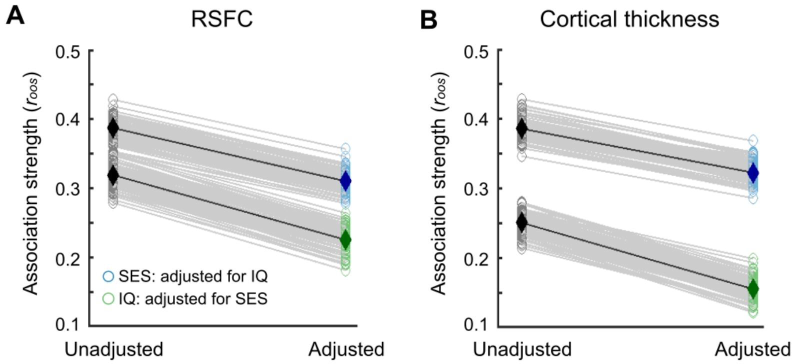


**fig. S17. Relative association strengths of SES vs. IQ scores.** Multivariate associations (out-of-sample correlations: $r_{oos}$) of **(A)** brain connectivity (RSFC) and **(B)** cortical thickness with IQ scores (green; NIH Toolbox Cognition Battery, total score) and SES (blue; social and economic domain of the Child Opportunity Index) using ridge regression (unadjusted: grey; left) in the Baseline ABCD sample (training sample: $n$ = 2,316; test sample: $n$ = 2,263). Multivariate out-of-sample brain-based associations of IQ scores were adjusted (see Methods) for SES (social and economic domain of the Child Opportunity Index; green; bottom right), while socioeconomics were adjusted by IQ scores (NIH Toolbox Cognition Battery, total score; blue; top right). Models were trained and tested on 100 split-half bootstrap samples. In all instances, the out-of-sample correlation ($r_{oos}$; correlation between predicted and observed scores) is plotted. Dark grey diamonds (left) represent the mean across 100 bootstrapped samples for unadjusted brain-based models of SES (top) and IQ scores (bottom). Dark blue and green diamonds (right) represent the mean across 100 bootstrapped samples for adjusted brain-based models of SES (blue; top) and IQ scores (green; bottom).

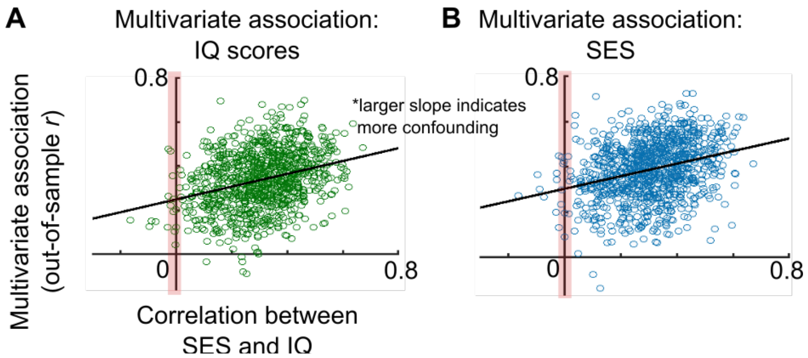


**fig. S18. Confound isolation cross-validation in multivariate brain-based association models.** Out-of-sample association (y-axis; out-of-sample correlation, $r_{oos}$) plotted against the correlation between SES (social and economic domain of the Child Opportunity Index) and IQ (NIH Toolbox Cognition Battery, total score) for **(A)** brain-based (resting-state functional connectivity (RSFC)) models (ridge regression) of IQ, treating SES as a confounding variable and for **(B)** brain-based (RSFC) models of SES, treating IQ scores as a confounding variable. The red shaded area in both panels represents the range in which SES and IQ scores are confounded (uncorrelated) in the test sample. The value in which the best fit line crosses through x = 0 (red shaded area) represents the out-of-sample association ($r_{oos}$) of brain-based models of IQ scores in panel **A** and SES scores in panel **B** in deconfounded test samples.

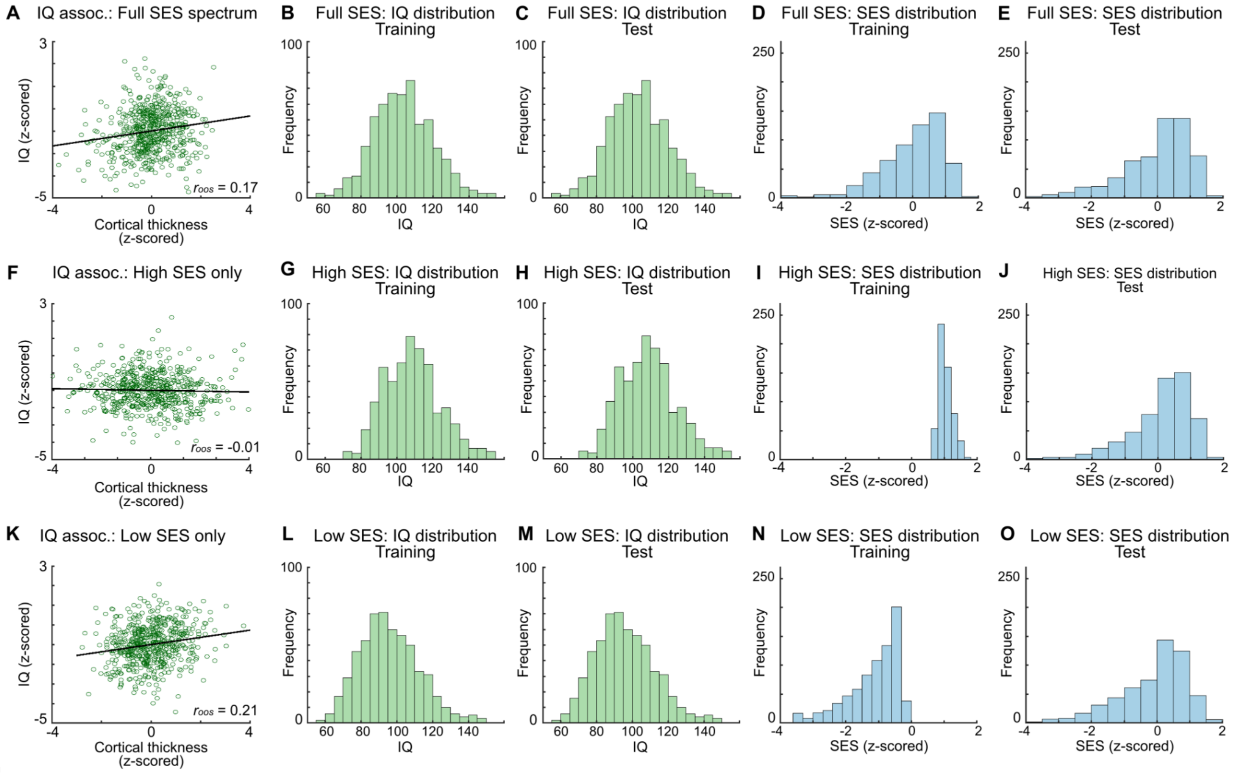


**fig. S19. Non-generalizability of multivariate brain-based associations of IQ scores. (A)** Out-of-sample IQ (NIH Toolbox Cognition Battery, all subscales) associations from cortical thickness using canonical correlation analysis (CCA), in the Adolescent Brain Cognitive Development (ABCD) Study. The y-axis shows the first canonical variates for IQ in the test data, scaled by the training weights. The x-axis shows the same but for cortical thickness. The correlation between the first canonical variates for IQ and cortical thickness is out-of-sample relative to the training data ($r_{oos}$). The training and test sets were subsampled from the pre-defined ABCD Discovery ($n$ = 2,351) and Replication ($n$ = 2,276) samples, respectively, thus covering the full ABCD SES spectrum (social and economic domain of the Child Opportunity Index) and matched for size ($n$ = 569). Subsample multivariate associations ($r_{oos}$ = 0.17; $n$ = 569) were lower than in the complete sample due to the known scaling of effect size with $n$. **(B)** IQ distribution in the training sample (full SES) **(C)** was matched to the test sample (full SES). This led to similar SES distributions in the **(D)** training and **(E)** test samples. **(F)** Out-of-sample IQ score with cortical thickness association, exactly as in **A**, but for a size-matched ($n$ = 569) training subsample drawn from only the high SES spectrum (z > 0.75). Restricting the training sample to high SES, while keeping the test sample as in **A**, (full SES) reduced multivariate association strength to $r_{oos}$ = -0.01 ($P$ = 0.49). **(G)** IQ distribution in the training sample (high SES) was matched to **(H)** the test sample (full SES). This restricted the **(I)** training SES distribution to high, **(J)** but not for the test sample. **(K)** Out-of-sample IQ score associated with cortical thickness association, exactly as in a, but for a size-matched ($n$ = 569) training subsample drawn from only the low SES spectrum (z < -0.55). Restricting the training sample to low SES, while keeping the test sample as in a (full SES) increased multivariate association strength to $r_{oos}$ = 0.21. **(L)** IQ distribution in the training sample (low SES) was matched to **(M)** the test sample (full SES). This restricted the **(N)** training SES distribution to low, **(O)** but not for the test sample.

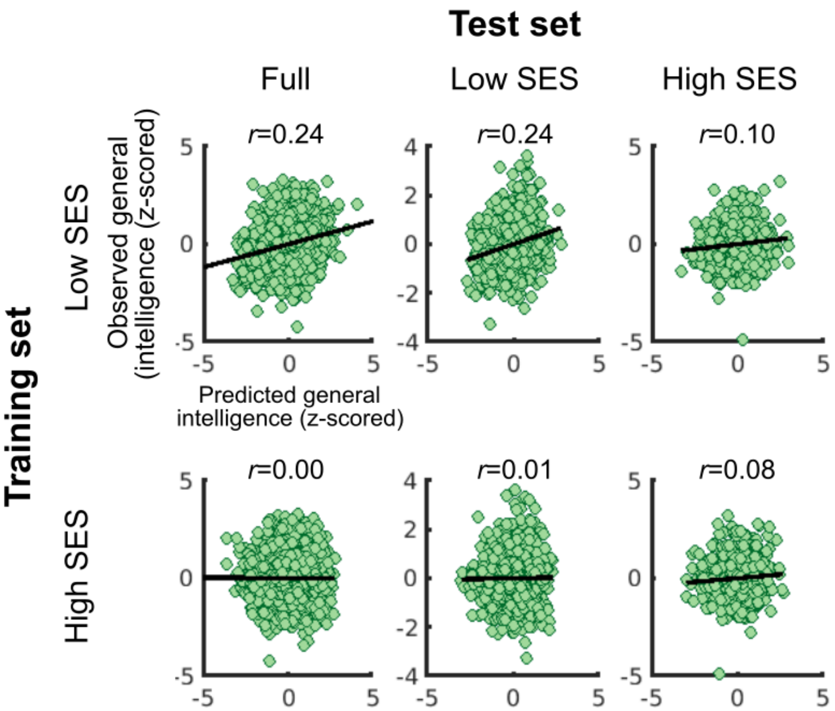


**fig. S20 | Generalizability of functional connectivity with IQ score associations using the connectome-based prediction model.** Out-of-sample (ABCD replication dataset) observed (y-axes) vs. predicted (x-axes) IQ scores with resting-state functional connectivity (RSFC) trained on individuals from low (*top row*) SES (normed Child Opportunity Index) backgrounds from the ABCD discovery dataset (low SES defined as less than 0.5 standard deviations from the median value [$z < -0.25$], where median $z = 0.25$) or individuals from high (*bottom row*) SES (normed COI $z > 0.75$) backgrounds and subsequently tested on an independent replication sample spanning the full replication set (*left column*), low SES (*middle row*), and high SES (*right row*) individuals.

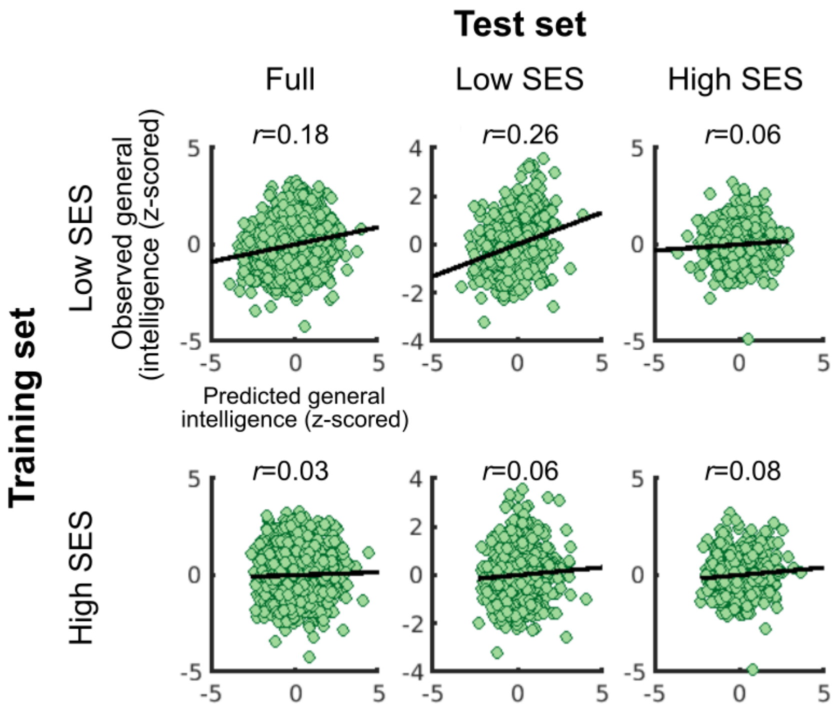


**fig. S21. Generalizability of cortical thickness with IQ score associations using the connectome-based prediction model.** Out-of-sample (ABCD replication dataset) observed (y-axes) vs. predicted (x-axes) IQ scores with cortical thickness trained on individuals from low (*top row*) SES (normed Child Opportunity Index) backgrounds from the ABCD discovery dataset (low SES defined as less than 0.5 standard deviations from the median value [$z < -0.25$], where median $z = 0.25$) or individuals from high (*bottom row*) SES (normed COI $z > 0.75$) backgrounds and subsequently tested on an independent replication sample spanning the full replication set (*left column*), low SES (*middle row*), and high SES (*right row*) individuals.

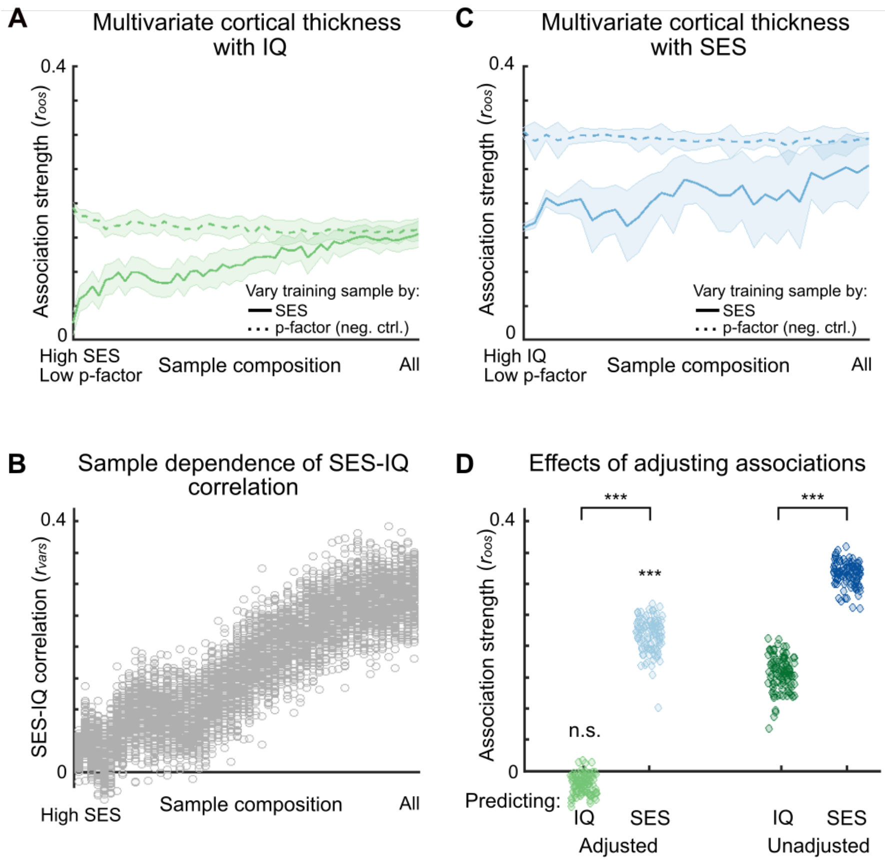


**fig. S22. Cross-contextual analyses test for shortcut learning in multivariate IQ and SES associations. (A)** IQ (NIH Toolbox Cognition Battery, all subscales) was predicted from cortical thickness using canonical correlation analysis (CCA), in the Adolescent Brain Cognitive Development (ABCD) Study (as in Fig. 6), with training data of varying compositions. The pre-defined ABCD Discovery (*n* = 2,316) data were repeatedly subsampled (*n* = 569) to range from distributions taken from the full dataset (x-axis, right) to restricted (left; high SES), based on SES (social and economic domain of the Child Opportunity Index; light blue), and psychopathology (p-factor; dark green; negative control). Multivariate associations, measured as out-of-sample correlation ($r_{oos}$) using the ABCD Replication sample (*n* = 2,263), are shown on the y-axis; line shading indicates one standard deviation (SD) around the mean $r_{oos}$ across 100 bootstrapped subsamples. **(B)** For the brain-based multivariate associations of IQ, varying by SES in **a**, the underlying correlations between SES and IQ ($r_{vars}$, y-axis; light blue), are shown as a function of SES composition of subsamples. **(C)** SES was predicted from cortical thickness, using CCA, in the ABCD (Baseline), with training data of varying SES compositions. The Discovery (*n* = 2,316) data were repeatedly subsampled (*n* = 569, 31 bins, 100 bootstraps per bin) to range from distributions from the full sample (x-axis, right) to restricted (left), based on IQ (light green; high IQ on left). Varying the training sample by psychopathology (p-factor; dark green), again served as a negative control. Multivariate associations ($r_{oos}$) are shown on the y-axis; line shading indicates one SD around the mean $r_{oos}$ across 100 bootstrapped samples. **(D)** Cortical thickness multivariate out-of-sample association of IQ fell to $r_{oos}$ = -0.02 (light green dots; IQ adjusted; $P = 0.62$, not significantly different from zero, mean across 100 bootstrapped samples) in subsamples in which there was a small correlation between SES and IQ ($r_{vars} < 0.10$; Fig. 6B). In subsamples with a stronger correlation between SES and IQ consistent with the ABCD sample ($r_{vars} \sim 0.30$), multivariate associations of IQ averaged $r_{oos}$ = 0.16 ($P < 0.001$; dark green dots; IQ unadjusted). Brain-based (cortical thickness) multivariate associations of SES remained robust at $r_{oos}$ = 0.22 (light blue dots; SES adjusted) in subsamples in which there was a small correlation between SES and IQ ($r_{vars} < 0.10$). In subsamples with a stronger correlation between SES and IQ consistent with the ABCD sample ($r_{vars} \sim 0.30$), cortical thickness out-of-sample associations of SES averaged $r_{oos}$ = 0.32 (dark blue dots; SES unadjusted).

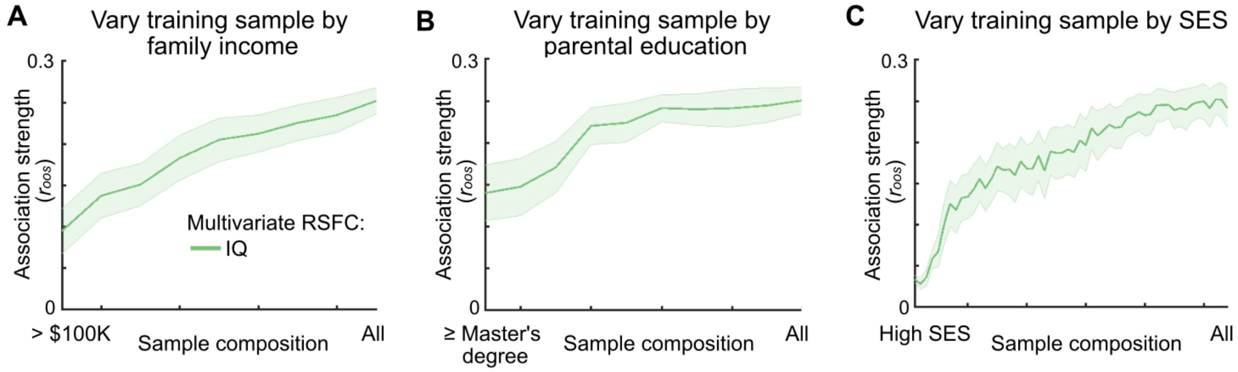


**fig. S23 | Effects of different socioeconomic measures on brain-based associations of IQ.** Multivariate (out-of-sample $r_{oos}$; y-axis; test set $n$ = 2,263) brain-based (resting-state functional connectivity (RSFC)) associations with IQ scores for varying levels of **(A)** family income, **(B)** parental education, and **(C)** SES (social and economic domain of the Child Opportunity Index) in the Baseline ABCD sample. For each socioeconomic variable, sample composition with respect to the socioeconomic variable of interest in the training (Discovery; $n$ = 569 in each bin) sample was varied, such that composition (x-axis) ranges from restricted (panel **A**: high income families (>100,000 USD); panel **b**: at least a Master's degree; panel **c**: individuals with high SES only) to maximum (subsamples from full ABCD sample training sample distribution). Note data plotted in panel **C** are from Fig. 6A.

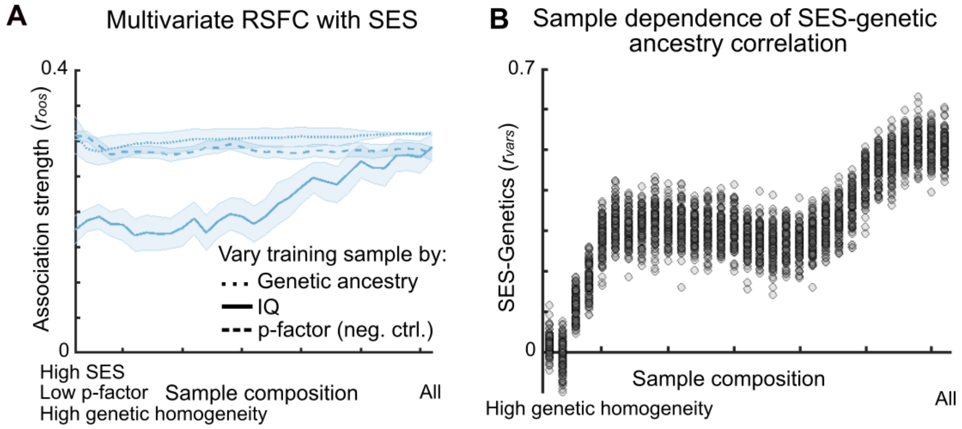


**fig. S24. Confound robustness of multivariate brain-based associations of SES. (A)** SES (all subscales of the social and economic domain of the Child Opportunity Index) were associated with resting-state functional connectivity (RSFC, 333 parcels), using canonical correlation analysis (CCA), in the Adolescent Brain Cognitive Development (ABCD) Study (as in Fig. 6C), with training data of varying sample compositions. The pre-defined ABCD Discovery ($n$ = 2,316) data were repeatedly subsampled ($n$ = 569) to range from a distribution from the full dataset (x-axis, right) to restricted (left; high IQ, low p-factor, more homogenous genetic ancestry). Multivariate associations, measured as out-of-sample correlation ($r_{oos}$) using the ABCD Replication sample ($n$ = 2,263), are shown on the y-axis; line shading indicates one standard deviation around the mean $r_{oos}$ across 100 bootstrapped subsamples. **(B)** Correlation between SES (social and economic domain of the Child Opportunity Index) and **(B)** the first genetic principal component (from ABCD 5.1 official release).

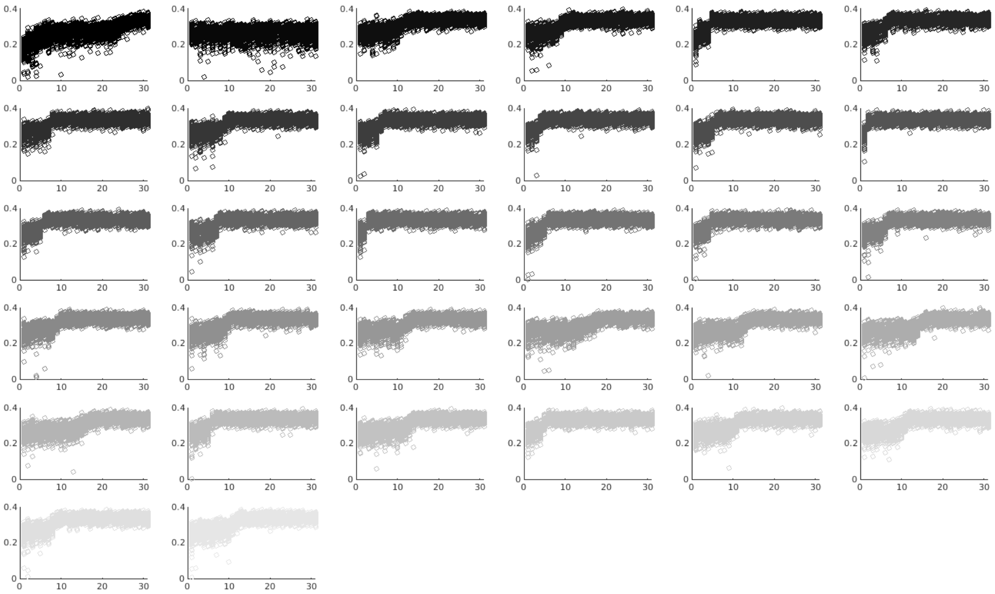

**fig. S25. Multivariate RSFC with SES associations across each genetic principal component.** SES (all subscales of the social and economic domain of the Child Opportunity Index) was predicted from resting-state functional connectivity (RSFC, 333 parcels), using canonical correlation analysis (CCA), in the Adolescent Brain Cognitive Development (ABCD) Study (as in Fig. 6, fig. S24), with training data of varying compositions with respect to each one of the 32 genetic principal components. The pre-defined ABCD Discovery (*n* = 2,316) data were repeatedly subsampled (*n* = 569) to range from sample compositions drawn from the full dataset (x-axis, right) to restricted with respect to genetic ancestry (left: more homogenous genetic ancestry). OPrediction accuracies, measured as out-of-sample correlation (y-axis; $r_{oos}$) using the ABCD Replication sample (*n* = 2,263) are shown on the y-axis. Each bootstrapped sample is plotted, represented by a single data point.

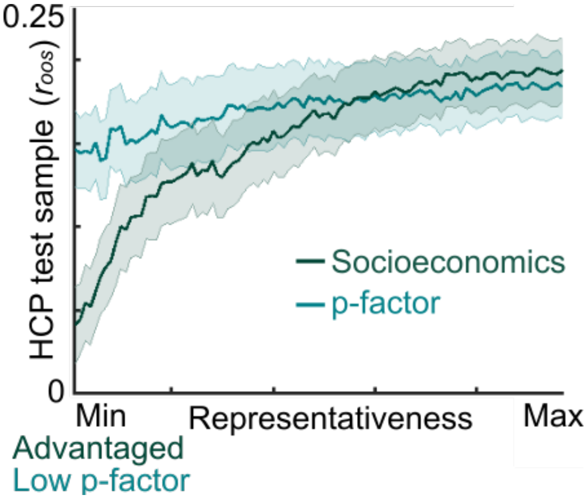


**fig. S26. Cross-sample generalizability of functional connectivity with IQ score associations.** Cross-sample (HCP) generalizability (out-of-sample $r$, $r_{oos}$; y-axis) of resting state functional connectivity (RSFC) with IQ scores (NIH Toolbox Cognition Battery subscales, all subscales) for varying SES compositions (x-axis; darker green) and total psychopathology (p-factor) composition (x-axis; lighter green) in the ABCD training set. All models were trained (CCA) using ABCD training (Discovery) data and tested out-of-sample in the adult HCP dataset (22-35 year olds). ABCD training sample composition (x-axis) ranges from restricted (SES: high SES individuals only; p-factor: no psychopathology) to all (subsample distributions drawn from full ABCD sample training sample). Each composition bin ($n$ = 97) was bootstrap resampled (with replacement) 100 times. Each bootstrapped ABCD training sample contained $n$ = 877 individuals to match the sample size of the HCP replication sample ($n$ = 877). The colored line represents the mean out-of-sample correlation for the first canonical vector in the HCP dataset. Shaded error bar represents one standard deviation (+/- 0.5) around the mean.

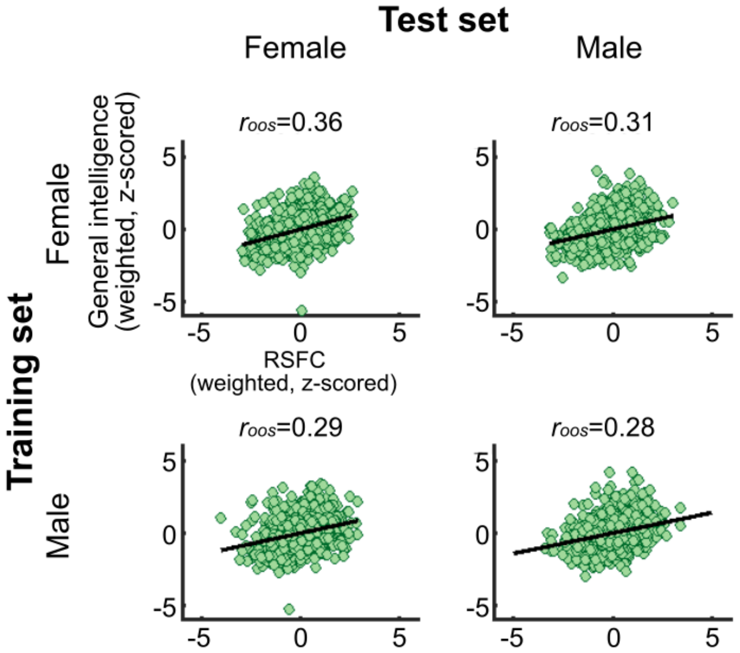


**fig. S27. Generalizability of functional connectivity associations with IQ across sexes.** Multivariate out-of-sample canonical correlations (first canonical vector; $r_{oos}$) for resting-state functional connectivity (RSFC) with IQ score (all subscales from the NIH Toolbox Cognition Battery) models trained on females (top row) from the ABCD sample or males (bottom row) and subsequently tested on an independent replication sample of females (left column) and males (right column). Unlike parametrically shifting socioeconomics in multivariate models of RSFC with general intelligence, RSFC with IQ models were generalizable across sex.

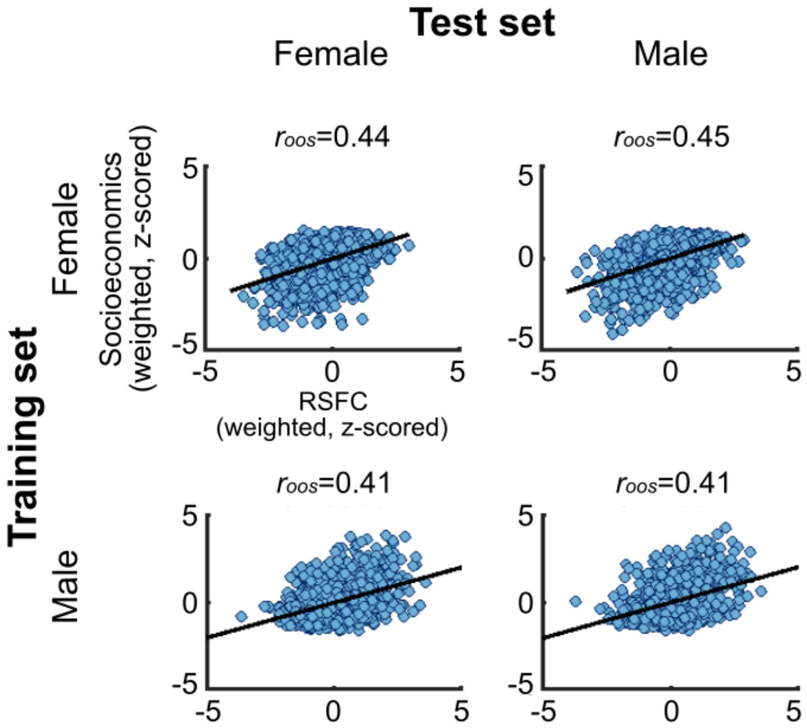


**fig. S28. Generalizability of functional connectivity associations with SES across sexes.** Multivariate out-of-sample canonical correlations (first canonical vector; $r_{oos}$) for resting-state functional connectivity (RSFC) with SES (social and economic domain of the Child Opportunity Index and family income) models trained on females (top row) from the ABCD sample or males (bottom row) and subsequently tested on an independent replication sample of females (left column) and males (right column). Like parametrically shifting general intelligence in multivariate models of RSFC with socioeconomics, RSFC with socioeconomics models were generalizable across sex.

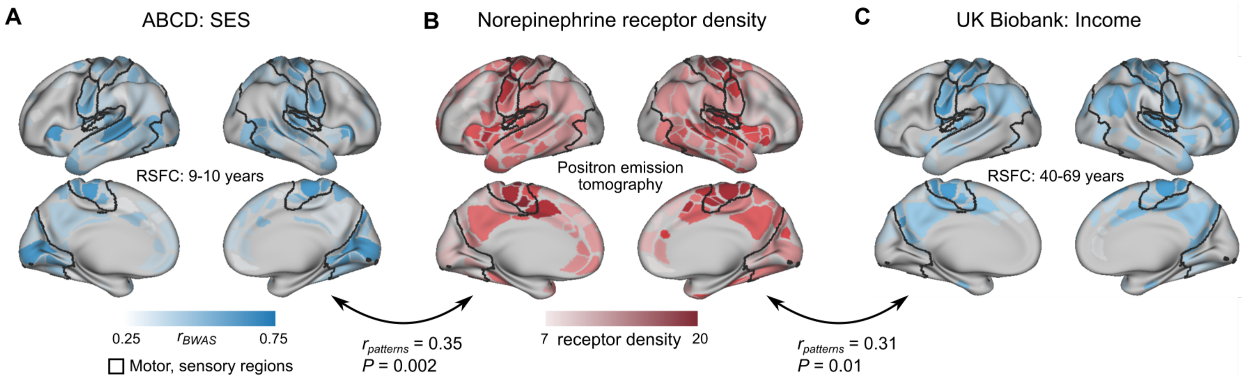


**fig. S29. Functional pattern comparisons against norepinephrine receptor density maps. (A)** Brain map of associations between resting-state functional connectivity (RSFC) and SES (social and economic domain of the Child Opportunity Index) in the baseline ABCD Discovery sample (*n* = 2,316, 9-10 years old). **(B)** Norepinephrine receptor density map from Positron Emission Tomography (PET) data (*45*) projected onto the 333 Gordon atlas (*117*). **(C)** Brain map of associations between RSFC and income in the UK Biobank dataset (*n* = 32,572, 40-69 years old). The correlation between brain maps and their associated *P*-value are shown between panels **A** and **B** and between panels **B** and **C**. Blue hues indicate relatively stronger associations in panels **A** and **C**. In panel **B**, red hues indicate greater norepinephrine receptor density.

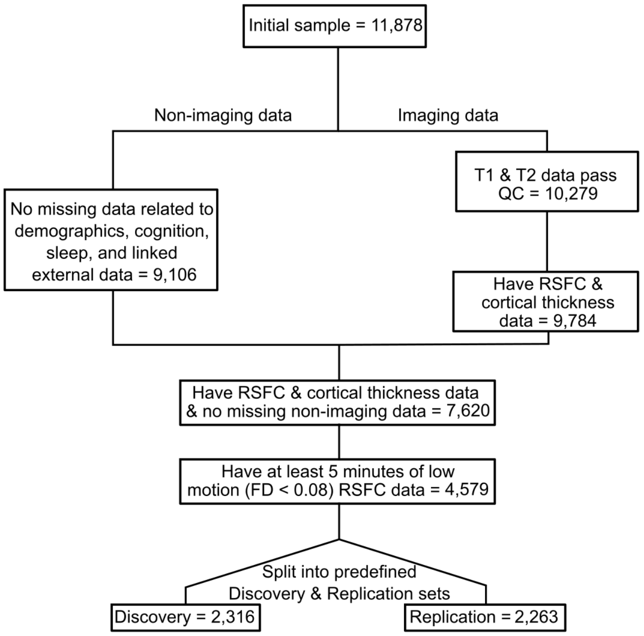


**Fig. S30. Consort diagram of participant flow for ABCD sample.** The diagram details how the sample was filtered from the initial Baseline Adolescent Brain Cognitive Development (ABCD) Study sample (*n* = 11,878). Imaging and non-imaging data were filtered for completeness and imaging data were further filtered based on data quality. Finally, the final sample was split into a Discovery sample (*n* = 2,316) and Replication sample (*n* = 2,263) that were matched across 9 variables (see Methods).
